# Ultrasensitive detection of circulating LINE-1 ORF1p as a specific multi-cancer biomarker

**DOI:** 10.1101/2023.01.25.525462

**Authors:** Martin S. Taylor, Connie Wu, Peter C. Fridy, Stephanie J. Zhang, Yasmeen Senussi, Justina C. Wolters, Wen-Chih Cheng, John Heaps, Bryant D. Miller, Kei Mori, Limor Cohen, Hua Jiang, Kelly R. Molloy, Brian T. Chait, Michael Goggins, Irun Bhan, Joseph W. Franses, Xiaoyu Yang, Mary-Ellen Taplin, Xinan Wang, David C. Christiani, Bruce E. Johnson, Matthew Meyerson, Ravindra Uppaluri, Ann Marie Egloff, Elyssa N. Denault, Laura M. Spring, Tian-Li Wang, Ie-Ming Shih, Euihye Jung, Kshitij S. Arora, Lawrence R. Zukerberg, Osman H. Yilmaz, Gary Chi, Bryanna L. Norden, Yuhui Song, Linda Nieman, Aparna R. Parikh, Matthew Strickland, Ryan B. Corcoran, Tomas Mustelin, George Eng, Ömer H. Yilmaz, Ursula A. Matulonis, Steven J. Skates, Bo R. Rueda, Ronny Drapkin, Samuel J. Klempner, Vikram Deshpande, David T. Ting, Michael P. Rout, John LaCava, David R. Walt, Kathleen H. Burns

## Abstract

Improved biomarkers are needed for early cancer detection, risk stratification, treatment selection, and monitoring treatment response. While proteins can be useful blood-based biomarkers, many have limited sensitivity or specificity for these applications. Long INterspersed Element-1 (LINE-1, L1) open reading frame 1 protein (ORF1p) is a transposable element protein overexpressed in carcinomas and high-risk precursors during carcinogenesis with negligible detectable expression in corresponding normal tissues, suggesting ORF1p could be a highly specific cancer biomarker. To explore the potential of ORF1p as a blood-based biomarker, we engineered ultrasensitive digital immunoassays that detect mid-attomolar (10^-17^ M) ORF1p concentrations in patient plasma samples across multiple cancers with high specificity. Plasma ORF1p shows promise for early detection of ovarian cancer, improves diagnostic performance in a multi-analyte panel, and provides early therapeutic response monitoring in gastric and esophageal cancers. Together, these observations nominate ORF1p as a multi-cancer biomarker with potential utility for disease detection and monitoring.

**Statement of Significance:** LINE-1 ORF1p transposon protein is pervasively expressed in many cancers and a highly specific biomarker of multiple common, lethal carcinomas and their high-risk precursors in tissue and blood. Ultrasensitive ORF1p assays from as little as 25 μL plasma are novel, rapid, cost-effective tools in cancer detection and monitoring.

## Introduction

There is significant clinical need for non-invasive methods to detect, risk stratify, and monitor cancers over time. Many malignancies are diagnosed at late stages when disease is widespread, contributing significantly to cancer morbidity and mortality(1). In contrast, there is a likely window in early-stage disease when patients are typically asymptomatic, in which treatments can be much more effective. Biomarkers are also needed to assess likelihood of progression in patients with precursor lesions, to provide prognostic information, and to predict and monitor responses or resistance to treatment(2). Considerable advances have been made towards detecting circulating tumor DNA, circulating tumor cells, microRNAs, and extracellular vesicles as non-invasive cancer biomarkers(3). However, achieving high sensitivities and specificities, particularly in affordable, scalable, clinical grade screening assays for early cancer detection, remains a major challenge. The plasma proteome provides a rich reservoir of potential biomarkers(4), which may be used individually or in combination for Multi-Cancer Early Detection (MCED) assays(5). However, most readily detectable proteins, including CA125 and HE4(6), FDA-cleared markers for the differential diagnosis of pelvic masses, are not sufficiently sensitive at the required high specificity(7) for cancer screening and/or are expressed in normal tissues and therefore lack the requisite specificity.

We have previously shown that expression of long interspersed element-1 (L1, LINE-1)-encoded open reading frame 1 protein (ORF1p) is a hallmark of many cancers(8), particularly p53-deficient epithelial cancers. These encompass many of the most commonly occurring and lethal human cancers, including esophageal, colorectal, lung, breast, prostate, ovarian, uterine, pancreatic, and head and neck cancers. L1 is the only active protein-coding transposon in humans. We each inherit, dispersed throughout our genomes, a complement of active L1 loci encoding two proteins: ORF1p, the highly expressed RNA binding protein(8), and ORF2p, an endonuclease and reverse transcriptase with limited expression(9) that generates L1 insertions in cancer genomes(10–13). L1 expression is repressed in normal somatic tissues, resulting in either very low or undetectable levels of L1 RNA and protein that appear to originate from epithelium(9,14). Epigenetic dysregulation of L1 and L1 ORF1p overexpression begin early in carcinogenesis, and histologic precursors of ovarian, esophageal, colorectal, and pancreatic cancers studied all express ORF1p at varying levels(8,15). ORF1p is thus a promising highly specific cancer biomarker.

Although elevated expression of ORF1p is readily detected by immunostaining in tumor tissue, ORF1p is found in plasma at low concentrations, well below detection limits of conventional clinical laboratory methods. We therefore applied the much more sensitive Single Molecule Arrays (Simoa), a digital bead-based ELISA technology, and in preliminary studies detected ORF1p in plasma at femtomolar levels in subsets of patients with advanced breast (33%, n=6)(16) and colorectal (90%, n=32)(17) cancers, respectively. Here, we assess the landscape of ORF1p plasma levels across multiple cancers, iteratively develop highly sensitive assays for potential applications in early or minimal residual disease detection, and provide evidence that plasma ORF1p may be an early indicator of therapeutic response.

## Results

Because our preliminary survey of plasma ORF1p levels by Simoa in patients with advanced stage colorectal cancer (CRC) indicated detectable ORF1p levels in 90% of cases(18), higher than the proportion of CRCs we previously reported to express ORF1p by immunohistochemistry (50%, n=18)(8), we first sought to benchmark ORF1p in tissues. Using a re-optimized protocol, we stained 211 CRCs [178 sequential cases included on a tissue microarray (TMA) as well as an additional 33 with matched plasma] and found 91% of CRC cases were immunoreactive for ORF1p (**Fig. 1a**). This result is consistent with genetic studies demonstrating somatic L1 retrotransposition in most CRCs(19), including activity in precancerous lesions antedating *APC* tumor suppressor loss(20–22). Similarly, genetic evidence shows esophageal adenocarcinoma (EAC) has high L1 activity(12), and L1 insertions occur in the highly prevalent Barrett’s esophagus (BE) precursor early in carcinogenesis(23,24). We therefore assembled a cross-sectional cohort of 72 BE cases with consensus diagnosis reached by three expert gastrointestinal pathologists. L1 RNA and ORF1p expression were pervasive in dysplastic BE and present in 100% of 51 esophageal carcinomas (**Fig. 1b,c**); all five BE cases indefinite for dysplasia and positive for ORF1p and/or L1 RNA developed high grade dysplasia on subsequent biopsies (not shown). Overall, this picture is similar to high grade serous ovarian cancers (HGSOC), where ORF1p is expressed in 90% of cases and 90% of fallopian tube precursor lesions (serous tubal intraepithelial carcinomas, STICs)(8,15,25). Taken together, ORF1p tissue expression is highly prevalent in gastrointestinal and gynecologic carcinomas and high-risk precursor lesions.

**Figure 1.**
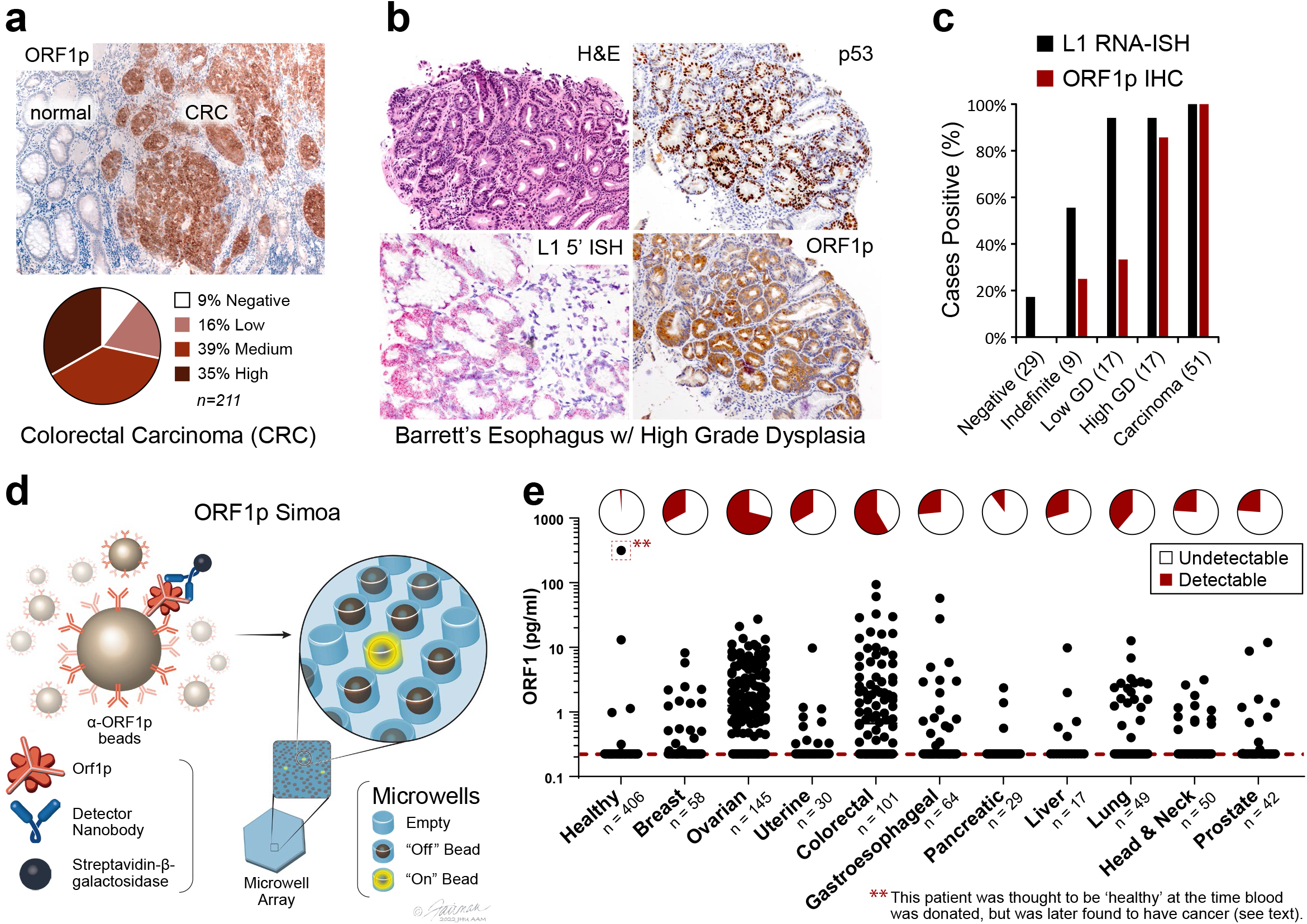
ORF1p expression is early and pervasive in carcinomas. **a**, ORF1p immunostaining in a cohort of 211 colorectal cancers. **b**, Representative BE case: lesional cells overexpress p53, the L1 RNA, and ORF1p. **c**, L1 RNA and ORF1p overexpression across a cohort of 72 consensus BE cases and 51 carcinomas. **d**, Schematic of single-molecule protein detection by Simoa; a second generation assay is shown. Antibody/nanobody-coated magnetic beads, present in excess relative to target, capture single target ORF1p molecules. Enzyme-labeled detection reagent (here, a homodimeric nanobody) is added, forming an “immunosandwich”, beads are loaded into microwells that each can hold at most one bead, and ORF1p molecules are then digitally detected using a fluorogenic substrate by counting “on” wells. First generation Simoa instead uses Nb5-coated beads and Ab6 detector. **e**, First-generation ORF1p Simoa detects plasma ORF1p with high specificity across major carcinomas. Pie charts indicate percentage of samples with detectable levels; dashed red line, LOD. **, this control patient was thought to be ‘healthy’ at the time blood was donated to the biobank but was later found to have prostate cancer and lymphoma.

We next sought to extend our tissue findings and explore plasma ORF1p. We optimized our previously reported ORF1p Simoa assay and assessed the landscape of ORF1p levels in pretreatment plasma from patients with advanced cancers. This “first-generation” assay uses a recombinant, single-domain camelid nanobody (Nb5) as the capture reagent and a monoclonal antibody (Ab6) as the detector reagent and has a limit of detection of 0.056 pg/mL (~470 aM trimeric ORF1p), corresponding to 1.9 fM in plasma after correcting for sample dilution (**Fig. 1d, Table S1**). With this assay, we surveyed multiple cancer types and >400 ‘healthy’ control individuals, who were without known cancer at the time blood was donated to the biobank. Plasma ORF1p appears to be a highly specific cancer biomarker, with undetectable levels in ~99% of controls (ages 20-90, **Fig. 1e, S1**). Of the five control patients with detectable ORF1p, the one with the highest ORF1p was later found to have advanced prostate cancer and a cutaneous T cell lymphoma; limited clinical information is available for the other four positive ‘healthy’ individuals. With a cutoff set at 98% specificity in healthy controls, the highest proportions of ORF1p(+) cases were observed in colorectal (58%, n=101) and ovarian cancers (71%, n=145). While most of these patients had advanced-stage disease, plasma ORF1p remained detectable in several early-stage patients in the cohort, including in those with ovarian and lung cancers and in 5/18 with intraductal papillary mucinous neoplasms in the pancreas (IPMN, **Fig. S2-S4**). Notably, four of eight stage I ovarian cancers in the cohort were positive (**Fig. S2**), suggesting that plasma ORF1p may be an indicator of early-stage disease. As L1 expression is also dysregulated in autoimmune disease and autoantibodies against ORF1p are prevalent in patients with systemic lupus erythematosus (SLE), we measured plasma ORF1p in 30 SLE patients and observed no detectable levels (**Fig. S5**)(26). Detectable ORF1p was seen in 1 of 30 patients with chronic liver disease; the one positive patient was subsequently diagnosed with hepatocellular carcinoma (**Fig. S5**). Size exclusion chromatography analysis of patient plasma further showed that the majority of ORF1p resides outside extracellular vesicles (**Fig. S6**). Together, these findings support the hypothesis that tumor-derived ORF1p can be found in the peripheral blood of cancer patients and may act as a cancer-specific biomarker.

Given the gap between proportions of ORF1p(+) cancers by tumor immunohistochemistry (~90% for CRC and HGSOC) versus by blood testing (~60-70%), we evaluated the possibility of increasing plasma assay sensitivity by decreasing the assay’s lower limit of detection. To this end, we developed a panel of ORF1p affinity reagents, including new recombinant rabbit monoclonal antibodies (RabMAbs) and engineered camelid nanobodies raised against recombinant human ORF1p. Because ORF1p is homotrimeric, we engineered multimeric nanobody reagents with the goal of enhancing binding affinity via increased avidity. These parallel development efforts ultimately yielded both improved nanobody and rabbit monoclonal antibody reagents with at least low-picomolar equilibrium dissociation constants (K_D_) (**Fig. S7-S12, Table S2-S4**). Iterative screening of these reagents with Simoa using recombinant antigen and select patient plasma samples yielded three best-performing capture::detection pairs, termed “second-generation,” which use rabbit monoclonal antibodies 34H7 and 62H12 as capture reagents and either Ab6 or homodimeric form of Nb5 (Nb5-5LL) as detector (**Fig. 2a-c, S13-S16**). Adding detergent further improved performance by limiting bead aggregation and improving bead loading into microwells. These second-generation assays comprised capture::detection pairs of 34H7::Nb5-5LL, 62H12::Nb5-5LL, and 62H12::Ab6, achieving detection limits of 0.016-0.029 pg/mL (130-240 aM trimeric ORF1p), and the four different reagents have predominantly non-overlapping epitopes in binning experiments (34H7 and 62H12 partially overlap, **Fig. 2a-c, Table S1, S5-S6**). Somewhat unexpectedly, analytical sensitivity did not perfectly correspond to clinical sensitivity. While the second-generation assays demonstrated less than an order-of-magnitude improvement in analytical sensitivity over the first-generation assay, they showed considerable improvement in circulating ORF1p detectability over background in buffer in re-measured samples across a large cohort of healthy and cancer patients (**Fig 2a, S17**). This difference may be due to differing accessibilities of circulating ORF1p epitopes or to different nonspecific binding patterns in plasma.

**Figure 2.**
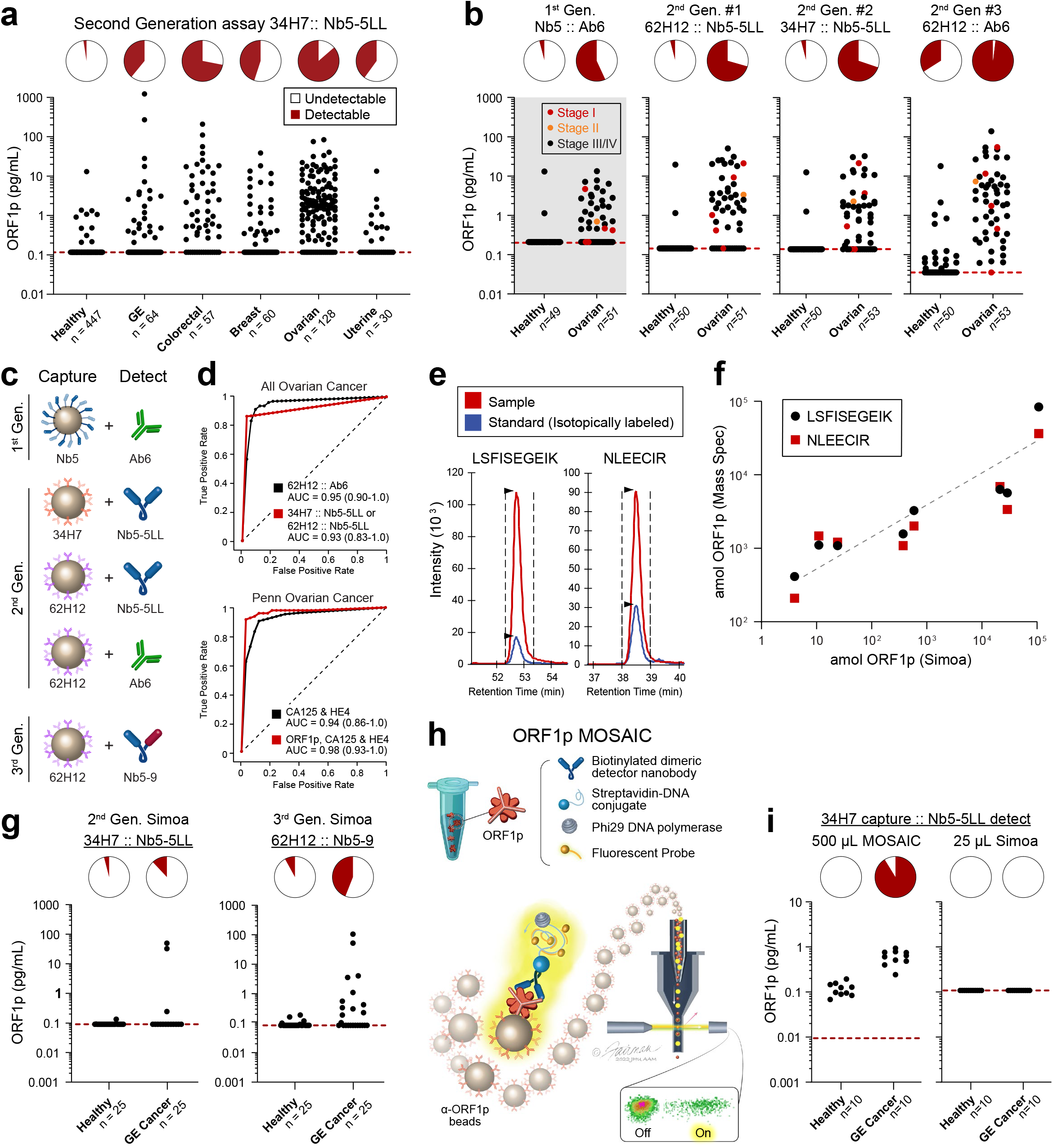
Improved detection of ORF1p with second- and third-generation assays. **a**, 34H7::Nb5-5LL second-generation assay measurements across a multi-cancer cohort. **b**, Ovarian cancer patients with age- and gender-matched controls in first- and second-generation assays; patients are a subset of those in 2a; red dots: stage I disease, orange dots: stage II disease. **c**, Schematic of affinity reagents used. 34H7 and 62H2 are custom mAbs; Nb5-5LL and Nb5-9 are an engineered homodimeric and heterodimeric nanobodies, respectively. **d**, ROC curves with single marker ORF1p across all healthy and ovarian cancer patients (top, n=128-132 cancer, 447-455 healthy), and multivariate models for ovarian (bottom, n=51-53 cancer, 50 healthy). **e**, Targeted proteomics measurements of plasma ORF1p from a gastric cancer patient using two quantotypic peptides (LSFISEGEIK and NLEECIR) with internal standards. **f**, correlation between measured ORF1p by Simoa and targeted proteomics assays; r=0.97 (Simoa vs LSFISEGEII) and r=0.99 (Simoa vs NLEECIR, t test), p<0.0001 for both. **g**, Comparison of 2^nd^ and 3^rd^ generation Simoa assays (25 μL) in 25 mostly undetectable gastroesophageal (GE) cancer and healthy control patients. **h**,Schematic of MOSAIC assays. Captured single molecule “immunosandwiches” are formed analogously to Simoa assays. DNA-conjugated streptavidin enables rolling circle amplification to be carried out, generating a strong local fluorescent signal on the bead surface, and then “on” and “off” beads are quantified by flow cytometry. **i**, 37H7::Nb5-5LL MOSAIC and Simoa assays in 10 previously-undetectable GE cancer and healthy control patients.

Undetectable or extremely low ORF1p levels in healthy individuals could readily be discriminated from measured ORF1p levels in ovarian cancer patients, resulting in a strong discriminatory ability with single-marker models (area under the receiver operating characteristic curve, AUCs of 0.93 to 0.948, sensitivity of 41% to 81% at 98% specificity, **Fig. 2d top panel, Table S7**). This large cohort included pre-treatment plasma samples from ovarian cancer patients (mostly high-grade serous ovarian carcinoma) with age-matched controls (n=51-53 women, **Fig 2b**); again, second-generation assays showed higher sensitivities while maintaining high specificities, notably achieving detection of five out of six Stage I/II patients at >98% specificity. Furthermore, multivariate models combining ORF1p (34H7::Nb5-5LL assay) with ovarian cancer biomarkers CA125 and HE4 yielded improved diagnostic performance over these existing markers (CA125 and HE4 alone, AUC = 0.94, 59% sensitivity at 98% specificity; ORF1p, CA125, and HE4, AUC = 0.98, 91% sensitivity at 98% specificity; **Fig 2d bottom panel, S18; Table S8**). While it is not clear whether the low ORF1p levels detected in several healthy individuals is due to nonspecific binding, true background levels of ORF1p, or an unappreciated pre-malignant state, several positive healthy controls were positive by only one of the three second-generation assays (n=4 positive by only 62H12::Nb5-5LL and n=75 positive by only 62H12:Ab6), suggesting nonspecific binding in at least some of these cases and the potential to improve specificity by combining data from multiple assays. Our results indicate that by developing improved affinity reagents, we achieved improved clinical sensitivity in detecting circulating ORF1p in cancer patients, with 83% sensitivity at >98% specificity towards early detection of ovarian cancer.

To further validate our results, we developed a targeted proteomics approach to measure ORF1p following affinity capture, with two distinct peptides measured vs. internal isotopically labeled control peptides (**Fig. 2e**). With this assay, we applied much larger volumes of plasma (3-6 ml, 120-240 fold more than the 25 μL used in Simoa assays) from a cohort of 10 patients, including 2 gastroesophageal (GE) cancer patients and one healthy control with very high ORF1p (230-1230 pg/ml), two healthy controls with high ORF1p, (3-5 pg/ml), and 5 healthy controls with low ORF1p (undetectable – 0.2 pg/ml). The results (**Fig. 2f, S19**) show strong correlation with Simoa, providing further confidence in our results (r=0.97-0.99, p<0.0001).

Building on the improvements made through nanobody engineering in our second-generation assays, we developed an expanded set of homodimeric, heterodimeric, and heterotrimeric anti-ORF1p nanobodies and screened them in combination with 34H7 and 62H12 capture antibodies, resulting in “third-generation” assays (**Figs. S9, S12, S20-21**). We noticed that reagents containing Nb2 performed very well in SPR but poorly in Simoa detection, and we hypothesized this was because Nb2 contains a lysine in the CDR, which would be biotinylated in the procedure, reducing affinity. We therefore engineered the new reagents to be C-terminally biotinylated on cysteine residues and varied linker sequence. Five of these assays, which utilize Nb2- and Nb9-containing constructs, outperform our second-generation assays in a cohort of 25 GE cancer patients with ORF1p measurements that were mostly undetectable previously, while maintaining high specificity versus healthy individuals (**Fig. 2g, S21**).

To leverage more sensitive assays for ORF1p detection, we next tested ORF1p affinity reagents from one of the second-generation Simoa assays on our recently developed Molecular On-bead Signal Amplification for Individual Counting platform (MOSAIC, **Fig. 2h**). MOSAIC develops localized on-bead signal from single captured molecules, in contrast to the microwell array format in Simoa, and improves analytical sensitivity by an order of magnitude over Simoa via increasing the number of beads counted(27). Furthermore, as the developed Simoa assays used only 25 μL plasma, we hypothesized that using larger plasma volumes would enhance ORF1p detectability by increasing the number of analyte molecules present. By using a 20-fold higher sample volume (500 μL plasma) and the MOSAIC platform, we achieved ten-fold higher analytical sensitivity, with a limit of detection of 0.002 pg/ml ORF1p (17 aM trimer, **Fig. S22**). Indeed, in a pilot cohort of gastroesophageal cancer and healthy patients, ORF1p levels in nine of ten previously undetectable cancer patients were readily discriminated from healthy individuals (**Fig. 2i**). Thus, in addition to improved affinity reagents, using larger sample volumes and more analytically sensitive technologies can further enhance both sensitivity and discrimination of circulating ORF1p levels between healthy controls and patients with cancer.

To test whether ORF1p might be useful for monitoring therapeutic response, 19 patients with gastroesophageal cancer were identified who had both detectable plasma ORF1p at diagnosis as well as subsequent samples available collected during or after treatment. Primary tumors were all adenocarcinoma and located in the esophagus (n=7), gastroesophageal junction (n=7) and stomach (n=5). All patients received systemic therapy. A smaller fraction of patients also received radiation and/or surgery (Supplement, **Table S9**). Clinical response (‘Responders’ and ‘Non-Responders’) was determined by review of re-staging CT and PET-CT imaging. Over an average of 465 days (range 98-1098), 12 patients died, six were alive at last follow-up (all ‘Responders’), and one was lost to follow-up. All 6 patients with detectable ORF1p at follow-up sampling, as defined by positivity over background in two of three assays, were also Non-Responders by imaging (**Fig. 3a**, p<0.0001, Fisher’s Exact test) and had reduced survival (p = 0.001 log-rank test for overall survival). In contrast, in all 13 Responders, circulating ORF1p dropped to undetectable levels post-treatment. Representative PET and PET-CT images are shown (**Fig. 3b**). Thus, reduction in circulating ORF1p paralleled treatment response and survival, while persistent circulating ORF1p corresponded to patients with refractory disease, indicating the predictive potential of this marker.

**Figure 3.**
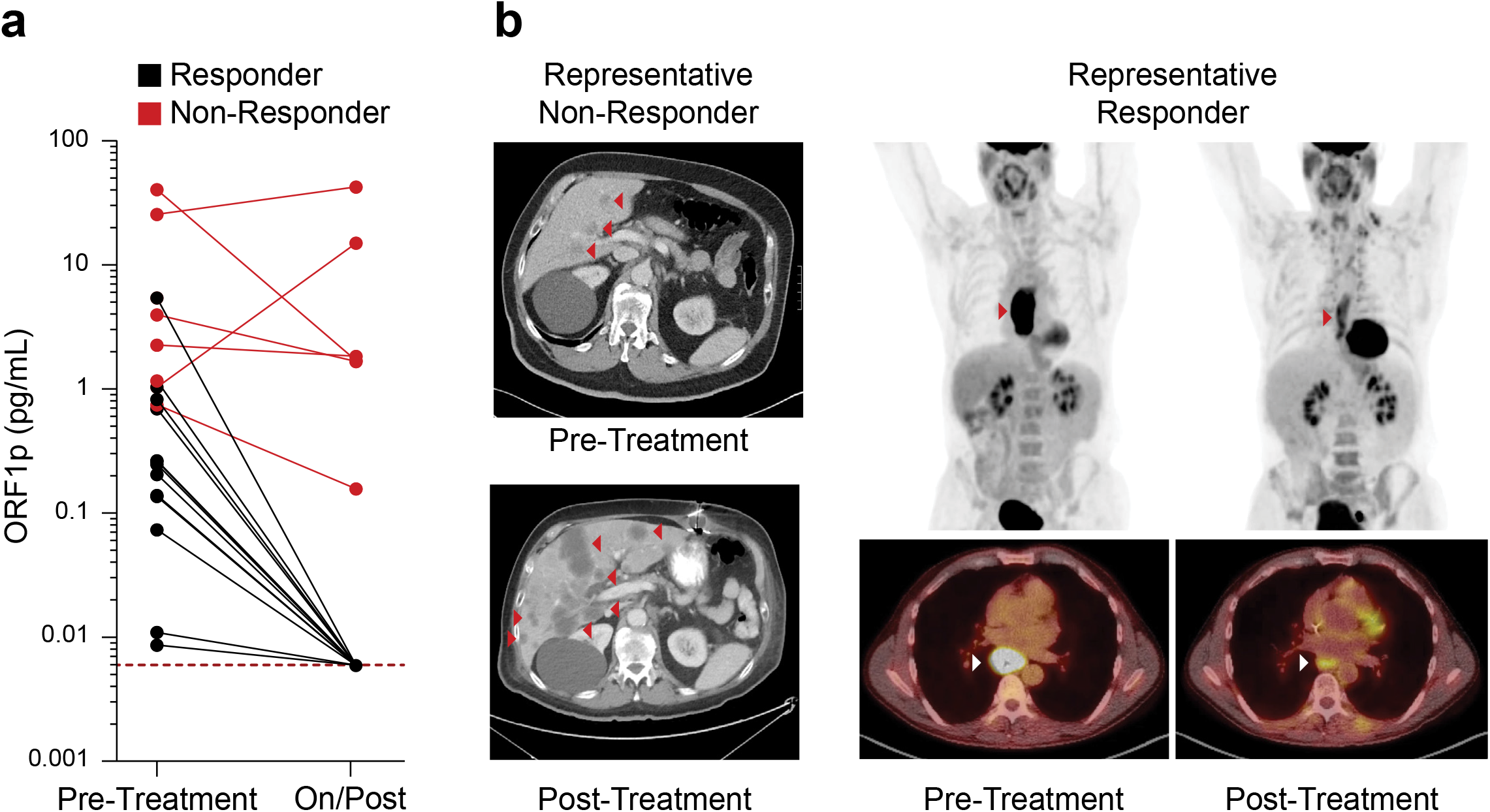
ORF1p is an early predictor of response in 19 gastroesophageal patients undergoing chemo/chemoradiotherapy. **a**, Plasma ORF1p as measured by all three second-generation Simoa assays before and during/post treatment; Responders and Non-Responders were characterized by post-therapy, pre-surgery imaging; p<0.0001, Fisher’s exact test. Non-Responders also have higher pre-treatment ORF1p than Responders (p=0.02, t-test). **b**, Representative CT and PET-CT from patients in the cohort.

## Discussion

Taken together, our data reveal for the first time that circulating ORF1p is a multi-cancer protein biomarker with potential utility across clinical paradigms, including early detection, risk stratification, and treatment response. These assays are enabled by ultrasensitive single-molecule detection technologies and high-quality affinity reagents, which are both required due to the attomolar-to-femtomolar circulating levels of ORF1p in cancer patients. Iterative improvements including optimized affinity reagents, buffer, and assay design yield highly sensitive and specific assays. A 20-fold volume scale-up to 500 μL appears promising for improving sensitivity without obviously compromising specificity, and this volume remains much smaller than a typical 5-10 mL blood draw and could be scaled further without limiting clinical applicability. The data strongly suggest that these assays are measuring *bona fide* tumor-derived circulating ORF1p for the following reasons: (1) four developed assays with predominantly non-overlapping high affinity reagents all measure similar levels across hundreds of samples; (2) levels appear specific to cancer patients, whose tumors overexpress ORF1p; (3) they correlate strongly with measurements made by targeted proteomics, and (4), plasma levels pre- and on/post treatment correlated with therapeutic response. Nonetheless, the low levels of circulating ORF1p makes orthogonal confirmation in larger cohorts by any other method challenging, as even the most sensitive mass spectrometry assays have limits of detection orders of magnitude higher.

The results expand our understanding that L1 expression is early and pervasive across carcinomas from multiple organs and high-risk precursor lesions, including dysplastic Barrett’s esophagus, which is challenging to diagnose and manage. Circulating ORF1p shows promise in early detection applications such as in ovarian cancer and may be more useful as part of a multi-analyte detection test combined with, for example, cfDNA methylation, longitudinal CA125 in ovarian cancer, or CEA in colorectal cancer(3,5,28). We demonstrate that ORF1p is an early indicator of chemotherapeutic response in gastric and esophageal cancers at timepoints where other parameters are often ambiguous, opening possibilities for monitoring minimal residual disease or relapse. Importantly, ORF1p appears to provide a level of specificity for cancers not achieved by other protein biomarkers, likely due to the unique biology of the retrotransposon, with repression of L1 in normal somatic tissue(9,13,14). ORF1p is therefore attractive as a putative “binary” cancer biomarker, in which a positive signal is highly specific for disease, with diagnostic utility both in tissue and plasma.

The assays are cost-effective (<$3 in consumables), rapid (<two hours), simple to perform, scalable, and have clinical-grade coefficients of variation (<15%). Flow cytometers for MOSAIC are common in clinical reference laboratories, and the assay could be modified for DNA-based readout by qPCR or sequencing. Limitations of the current work include the relatively small numbers of early-stage samples and a small and heterogeneous gastroesophageal therapeutic cohort. Larger cohorts will be needed for further validation. Further optimizations to both assay design and reagents will likely be possible, and larger cohorts are needed to further validate and develop third generation Simoa assays and MOSAIC assays. Finally, it is unclear how ORF1p, which is normally cytosolic, enters the blood and what clinicopathologic factors might affect these levels. Future work will also be needed to understand whether there is a normal baseline level of circulating ORF1p, as implied by the trace amounts seen when ORF1p was measured from much larger volumes of plasma using targeted mass spectrometry, and what factors affect this level.

## Methods

Provided in detail in *Supplementary Information.*

## Supporting information

Merged Supplement

## Acknowledgements

We thank Jeni Fairman for illustrations and Bert Vogelstein for plasma samples from colorectal cancer patients. We are grateful to Phil Cole for resources for protein expression and purification and helpful discussions and to Andrew Kruse and Edward Harvey for helpful discussions regarding nanobodies. We thank Zuzana Tothova for helpful discussion and review of the manuscript. This work was supported by the National Institutes of Health grants R01GM130680 (KHB), K08DK129824 (MST), F32EB029777 (CW), R01CA240924 (DTT), U01CA228963 (DTT), P41 GM109824 (MPR, BTC), T32CA009216 (MST, GE), U01CA233364, U2CCA271871, U01CA152990 (SJS), R01GM126170 (JL), P50CA228991 Ovarian SPORE (EJ, T-LW, I-MS, RD); Break *Through* Cancer (KHB); Earlier.Org (KHB and DRW); Minnesota Ovarian Cancer Alliance (KHB); DOD W81XWH-22-1-0852 (EJ, RD); Canary Foundation (RD); Gray Foundation (EJ, RD); The Concord (MA) Detect Ovarian Cancer Early Fund (SJS), Good Ventures (Open Philanthropy Project); Friends of Dana-Farber Cancer Institute; Dana-Farber Cancer Institute; and the Dana-Farber/Harvard Cancer Center (DF/HCC); ACD-Biotechne (DTT, VD); Robert L. Fine Cancer Research Foundation (DTT); Worldwide Cancer Research grant 19-0223 (JL); Robertson Therapeutic Development Fund (JL); Nile Albright Research Foundation (BRR); Vincent Memorial Research Foundation (BRR); SU2C Gastric Cancer Interception Research Team Grant (SU2C-AACR-DT-30-20, SJK, DTT, administered by the American Association for Cancer Research, the Scientific Partner of SU2C).

## Author Contributions

MST, CW, ÖHY, SJK, VD, DTT, JL, DRW, and KHB formulated the research plan and interpreted experimental results with assistance from SJZ, LC, YS, JCW, WCC, JH, BDM, and HJ. CW, SZJ, LC, and YS performed Simoa and MOSAIC experiments. WCC, JH, HJ, and BDM performed biochemical experiments. GE performed mouse experiments and interpreted results. PCF, MST, CW, HJ, KRM, BTC, MPR, and JL developed and engineered nanobody constructs. PCF performed SPR affinity measurements. MST, BDM, JCW, and JL designed and performed mass spectrometry experiments. MG, IB, JWF, XY, MET, XW, DC, BEJ, MM, RU, AME, END, LMS, TLW, IMS, EJ, BV, GC, BLN, ARP, MS, UAM, BRR, RD, SJK, and DTT provided patient samples and data and interpreted clinical results. SJS and KM carried out bioinformatic analysis. MST, LRZ, OHY, and VD diagnosed biopsies, scored cases, and interpreted results. MST, CW, DRW, and KHB wrote the manuscript. All authors edited and approved the manuscript.

