## Supplementary material for "Ultrasensitive detection of circulating LINE-1 ORF1p as a specific multi-cancer biomarker": Merged Supplement

**Supplementary Information**

**Materials and Methods**

**Materials.** All affinity reagents used in this work are listed in the Supplementary Information (Table S2). Conjugation reagents, paramagnetic beads, and assay buffers were obtained from Quanterix Corporation. DNA oligos used in the MOSAIC assay were obtained from Integrated DNA Technologies. Antibodies used in final Simoa and MOSAIC assays (monoclonals Ab6, Ab54, 62H12, 34H7) were additionally validated by western blotting (**Figure S26**).

**Preparation of capture and detector reagents.** All capture antibodies and nanobodies were obtained in or dialyzed into phosphate buffered saline (PBS). For the first-generation Simoa assay, $7\times10^8$ carboxylated paramagnetic 2.7-$\mu$m beads (Homebrew Singleplex Beads, Quanterix Corp.) were first washed three times with 400 $\mu$L Bead Wash Buffer (Quanterix Corp.) and two times with 400 $\mu$L cold Bead Conjugation Buffer (Quanterix Corp.) before being resuspended in 390 $\mu$L cold Bead Conjugation Buffer. A 1 mg vial of 1-ethyl-3-(3-dimethylaminopropyl) carbodiimide hydrochloride (EDC) (Thermo Fisher Scientific) was then dissolved to 10 mg/mL in cold Bead Conjugation Buffer, and 10 $\mu$L was added to the beads. The beads were shaken for 30 minutes at 4°C to activate the carboxyl groups on the beads, which were then washed once with 400 $\mu$L cold Bead Conjugation Buffer and resuspended in the capture nanobody solution (10 $\mu$g nanobody total), diluted in Bead Conjugation Buffer to a final volume of 400 $\mu$L. The beads were shaken for two hours at 4°C, washed twice with 400 $\mu$L Bead Wash Buffer, and resuspended in 400 $\mu$L Bead Blocking Buffer (Quanterix Corp.) before shaking at room temperature for 30 minutes to block the beads. After one wash each with 400 $\mu$L Bead Wash Buffer and Bead Diluent (Quanterix Corp.), the beads were resuspended in Bead Diluent and stored at 4°C. Beads were counted with a Beckman Counter Z Series Particle Counter before using in assays. For second-generation Simoa assays, the following bead coupling conditions were used: $4.2\times10^8$ starting beads, 300 $\mu$L wash volumes, 6 $\mu$L EDC, and 40 $\mu$g antibody.

For biotinylation of detector antibodies or nanobodies, a 1 mg vial of Sulfo-NHS-LC-LC-biotin was freshly dissolved in 150 $\mu$L water and added at 80-fold molar excess to a 1 mg/mL solution of antibody or nanobody. The reaction mixture was incubated at 30 minutes at room temperature and subsequently purified with an Amicon Ultra-0.5 mL centrifugal filter (50K and 10K cutoffs for antibody and dimeric nanobody, respectively). Five centrifugation cycles of 14,000xg for five minutes were performed, with addition of 450 $\mu$L PBS each cycle. The purified biotinylated detector reagent was recovered by inverting the filter into a new tube and centrifuging at 1000xg for two minutes. Concentration was quantified using a NanoDrop spectrophotometer.

**Recombinant ORF1p protein production.** ORF1p was prepared as described(1); briefly, codon optimized human ORF1p corresponding to L1RP (L1 insertion in X-linked retinitis pigmentosa locus, GenBank AF148856.1) with N-terminal His6-TEV was expressed in E. Coli, purified by Ni-NTA affinity, eluted, tag cleaved in the presence of RNaseA, and polished by size exclusion in a buffer containing 50 mM HEPES pH 7.8, 500 mM NaCl, 10 mM MgCl2, and 0.5 mM tris(2-carboxyethyl) phosphine (TCEP), resulting in monodisperse trimeric ORF1p bearing an N-terminal glycine scar.

**Nanobody generation and screening.** Nanobodies were generated essentially as decribed(2,3) using mass spectrometry/lymphocyte cDNA sequencing to identify antigen-specific nanobody candidates. Briefly, a llama was immunized with monodisperse ORF1p, and serum and bone marrow were isolated. The heavy chain only IgG fraction (VHH) was isolated from serum and bound to a column of immobilized ORF1p. Bound protein was eluted in SDS and sequenced by mass spectrometry, utilizing a library derived from sequencing VHH fragments PCR-amplified from bone marrow-derived plasma cells. Candidate sequences were cloned into an E. coli expression vector with C-terminal His6 tag and expressed in 50 ml cultures in E. coli Arctic Express RP (Agilent) with 0.2 mM IPTG induction at 12°C overnight. Periplasmic extract was generated as follows: pellets were resuspended in 10 ml per L culture TES buffer (200 mM Tris-HCl, pH 8.0, 0.5 mM EDTA, and 500 mM sucrose), 20 ml/L hypotonic lysis buffer added (TES buffer diluted 1:4 with ddH2O), supplemented with 1 mM PMSF, 3 µg / ml Pepstatin A, incubated 45 min at 4°C, and centrifuged at 25,000 x g for 30 min. The supernatant (periplasmic extract) was bound to ORF1p-conjugated Sepharose, washed 3 times, eluted with SDS at 70°C for 10 min, and periplasmic extract and elution were analyzed by SDS-PAGE to assay expression and yield. ORF1p-binding candidates were purified as below and analyzed by ELISA (**Figure S7**).

**Nanobody and multimeric nanobody purification.** C-terminally His6-tagged nanobody constructs were expressed and purified essentially as described(2). Briefly, protein was expressed in E. coli Arctic Express RP (Agilent) with 0.2 mM IPTG induction at 12°C overnight. Periplasmic extract (generated as above) was supplemented with 5 mM $MgCl_2$, 500 mM NaCl, and 20 mM imidazole, purified by Ni-NTA chromatography, dialyzed into 150 mM NaCl, 10 mM HEPES, pH 7.4, and concentrated to 1-3 mg/ml by ultrafiltration. “5xCys tail” constructs were purified with the addition of 5 mM TCEP-HCl in resuspension, wash, elution, and dialysis buffers.

**Surface plasmon resonance (SPR) assays.** Binding kinetics ($k_a$, $k_d$, and $K_D$**)** of antibody and nanobody constructs for ORF1p were obtained on a Biacore 8K instrument (Cytiva). Recombinant ORF1p was immobilized on a Series S CM5 sensor chip at 1.5 µg/ml using EDC/NHS coupling chemistry according to the manufacturer’s guidelines. Nanobodies and antibodies were prepared as analytes and run in buffer containing 20 mM HEPES pH 7.4, 150 mM NaCl, and 0.05% Tween-20. Analytes were injected at 30 µl/min in single-cycle kinetics experiments at concentrations of 0.1, 0.3, 1, 3.3, and 10 nM, with association times of 120-180 sec, and a dissociation time of 1200-7200 sec, depending on observed off-rate. Residual bound protein was removed between experiments using 10 mM glycine-HCl pH 3.0. Data were analyzed using Biacore software, fitting a Langmuir 1:1 binding model to sensorgrams to calculate kinetic parameters.

For epitope binning, pairs of antibodies were sequentially flowed over immobilized ORF1p using Biacore tandem dual injections according to the manufacturer’s guidelines. Antibodies were injected at concentrations of 200 nM with a flow rate of 10 µl/min. Contact time for the first antibody was 120 sec, followed by 150 sec for the second antibody, then a 30 sec dissociation. Response signal for the second antibody was measured in a 10 sec window at the beginning of dissociation. The chip was regenerated between experiments with glycine pH 3.0 as above. Data were analyzed using the Biacore software epitope binning module.

**ORF1p Simoa assays.** Simoa assays were performed on an HD-X Analyzer (Quanterix Corp.), with all assay reagents and consumables loaded onto the instrument according to the manufacturer’s instructions. 250,000 capture beads and 250,000 helper (non-conjugated) beads were used in each Simoa assay. A three-step assay configuration was used for the first- and second-generation assays, consisting of a 15-minute target capture step (incubation of capture beads with 100 $\mu$L sample), 5-minute incubation with detector reagent (0.3 $\mu$g/mL for both first- and second-generation assays), and 5-minute incubation with streptavidin-$\beta$-galactosidase (150 pM for first-generation assay; 300 pM for second-generation assays). The beads were washed with System Wash Buffer 1 (Quanterix Corp.) after each assay step. Upon the final wash cycle, the beads were loaded together with the fluorogenic enzyme substrate resorufin $\beta$-D-galactopyranoside into a 216,000-microwell array, which was subsequently sealed with oil. Automated imaging and counting of “on” and “off” wells and calculation of average enzyme per bead (AEB) were performed by the instrument. Calibration curves were fit using a 4PL fit with a $1/y^2$ weighting factor, and the limit of detection (LOD) was determined as three standard deviations above the blank.

All plasma and serum samples were diluted four-fold in Homebrew Sample Diluent (Quanterix Corp.) with 1x Halt Protease Inhibitor Cocktail (ThermoFisher), with an additional 1% Triton-X 100 added in the second-generation assays. All recombinant ORF1p calibrators were run in triplicates, with four replicates for the blank calibrator, and all plasma and serum samples were run in duplicates. The average LOD across all sample runs was determined for each assay and depicted in each figure.

Healthy individual plasma and serum samples were obtained from the Mass General Brigham Biobank, with additional samples from the Penn Ovarian Cancer Research Center and Tomas Mustelin (University of Washington). Additional breakdown of patients within each cancer type, by demographic and clinicopathological variables, where available, is included in **Figures S2-S3, and S19-21, and Table S10**.

**ORF1p large-volume MOSAIC assays.** MOSAIC assays were performed as previously described, using 2 ml microcentrifuge tubes for the initial capture step(4). For each sample, 500 $\mu$L plasma was diluted four-fold in

Homebrew Sample Diluent with protease inhibitor and 1% Triton-X 100 to a total volume of 2 mL. Briefly, 100,000 capture beads were incubated with sample and mixed for two hours at room temperature, followed by magnetic separation and resuspended in 250 $\mu$L System Wash Buffer 1 before transferring to a 96-well plate. The beads were then washed with System Wash Buffer 1 using a Biotek 405 TS Microplate Washer before adding 100 $\mu$L nanobody detector reagent (0.3 $\mu$g/mL, diluted in Homebrew Sample Diluent) and shaking the plate for 10 minutes at room temperature. After washing with the microplate washer, the beads were incubated with 100 $\mu$L streptavidin-DNA (100 pM, diluted in Homebrew Sample Diluent with 5 mM EDTA and 0.02 mg/mL heparin) with shaking for 10 minutes at room temperature, followed by another washing step. The beads were transferred to a new 96-well plate, manually washed with 180 $\mu$L System Wash Buffer 1, and resuspended in 50 $\mu$L reaction mixture for rolling circle amplification (RCA). The RCA reaction mixture consisted of 0.33 U/uL phi29 polymerase, 1 nM ATTO647N-labeled DNA probe, 0.5 mM deoxyribonucleotide mix, 0.2 mg/mL bovine serum albumin, and 0.1% Tween-20 in 50 mM Tris-HCl (pH 7.5), 10 mM $(NH_4)_2SO_4$, and 10 mM $MgCl_2$. The beads were shaken at 37°C for one hour, followed by addition of 160 $\mu$L PBS with 5 mM EDTA and 0.1% Tween-20. After washing the beads once with 200 $\mu$L of the same buffer, the beads were resuspended in 140 $\mu$L buffer with 0.2% BSA. All samples were analyzed using a NovoCyte flow cytometer (Agilent) equipped with three lasers. Analysis of average molecule per bead (AMB) values was performed as previously described using FlowJo software (BD Biosciences) and Python. All code used for MOSAIC data analysis can be downloaded as part of the waltlabtools.mosaic Python module, which is available at https://github.com/tylerdougan/waltlabtools.

**Targeted proteomics analysis of immunoprecipitated ORF1p.** Protein levels the LINE-1 ORF1p (UniProt ID: Q9UN81) were determined with targeted proteomics using isotopically-labeled standard peptides (AQUA QuantProHeavy peptides with $^{13}C^{15}N$-labeled C-terminal lysine or arginine, Thermo Fisher) for accurate quantification. Assays were developed for two quantotypic peptides of ORF1p, namely LSFISEGEIK and cysteine-alkylated NLEECITR (manuscript in preparation,  the approach is similar to the assay development described previously for other proteins(5)). Briefly, 3-6 mL patient plasma was diluted with an equal volume of 2x dilution buffer (PBS containing 2% Triton X-100, 10 mM EDTA, and 1 Pierce protease inhibitor tablet per 25 ml (2x concentration, Thermo) for a final concentration of 1% Triton X-100, 5 mM EDTA, and 1x protease inhibitor and bound to 7 million 62H12-conjugated magnetic beads for 1 hour at room temperature. Beads were washed 3 times with 5x PBS containing 0.1% tween 20 and 1x protease inhibitor, then once with the same buffer lacking tween 20, and eluted in 50 µl buffer containing 2% SDS and 50 mM Tris pH 8.5 by heating for 5 minutes at 95°C with agitation. Separated eluates were subjected to in-gel digestion using trypsin (150 ng sequencing grade modified trypsin V5111; Promega) after reduction with 10 mmol/L dithiothreitol and alkylation with 55 mmol/L iodoacetamide proteins, prior to LC-MS analyses of the target peptides(5).

**Classification models.** Classification models were trained for (1) all healthy and all ovarian cancer patients measured by the second-generation assays; and (2) the subset of 51 ovarian cancer and 50 age-matched healthy female patients, obtained from Ronny Drapkin (University of Pennsylvania). Each dataset contained no missing values, and the measurements in the datasets were log-transformed and normalized beforehand for classification analysis of healthy and ovarian cancer subjects. Logistic regression was used for the univariate classifier and the k-nearest neighbors (KNN) and light gradient-boosting machine (LightGBM), which had the best performances among the classifiers, were used for the multivariate classifier, and implemented in Python 3.7.15 with scikit-learn version 1.0.2 package. Each classifier was given a weight optimization between classes to deal with data imbalance between healthy and cancer subjects, as well as hyperparameter tuning using grid search.

The performance of each biomarker in differentiating ovarian cancer subjects from healthy subjects was evaluated with fivefold cross validation by calculating accuracy, precision, recall, f1-value, sensitivity, specificity, and area under the receiver-operating characteristic (ROC) curve (AUC). A stratified five-fold cross-validation strategy randomly splits the positive and negative samples into five equally sized subsets. One positive subset and one negative subset were selected as the test dataset each time, and the other samples were used to train a classification model.

In the multivariate analysis, the Variance Inflation Factor (VIF) for the biomarkers was calculated, and any biomarkers with extremely high correlation with VIF greater than 10 were excluded from the classification model in advance.

**Barrett's esophagus cases.** A cohort of 75 esophageal biopsies with BE and varying degrees of dysplasia were assembled. Negative cases were screened to have no prior history of dysplasia. The mean age of the cohort was 67 years with a male predominance (M:F ratio = 3.7:1). All samples were re-analyzed for histological features of dysplasia by three experienced gastrointestinal pathologists (LRZ, VD, OHY) who were blinded to the original diagnosis. A consensus was reached for 72 cases and the consensus diagnosis was used as the gold standard. There was moderate agreement between pathologists (kappa 0.43-0.51).

**Colon cancer tissue microarray**. 178 sequential CRCs resected by a single surgeon from 2011-2013 were assembled on a 3 mm core tissue microarray. All cases were independently scored by two pathologists. The mean age of the cohort was 65 years with 49.8% males. Mean follow-up was 25 months. At resection, 23% were stage I, 33% were stage II, 44% were stage III, and 1% were stage IV.

**Ovarian Cancer Samples**. Age-matched ovarian cancer (n=53) and healthy control (n=50) patient plasma samples were from University of Pennsylvania Ovarian Cancer Research Center, OCRC Tumor BioTrust Collection, Research Resource Identifier (RRID): SCR_02287.

**Gastroesophageal cancer treatment cohort.** Nineteen patients received systemic therapy, 3 of which also underwent surgical resection. Patients were treated with concurrent chemotherapy (carboplatin/taxol) and radiation (N=3), fluorouracil/ leucovorin/ oxaliplatin/ docetaxel (FLOT, N=2), fluorouracil/ leucovorin/ irinotecan/ oxaliplatin (FOLFIRINOX, N=2), fluorouracil/ leucovorin/ oxaliplatin (FOLFOX, N=9), FOLFOX + trastuzumab (N=1), pembrolizumab (N=1) or FOLFOX then chemoradiation (1). The mean age of the cohort was 76 years. All patients were male (100%). Fifty-eight percent had locally advanced disease (stage II-III) and 42% had advanced disease (stage IV) at the time of initial diagnosis. Sixty-eight percent (N=13) were deemed responders to therapy while 32% (N=6) were deemed non-responders to standard therapy on review of re-staging imaging (CT and/or PET-CT) by investigators blinded to the assay results. Note that the on/post-treatment blood draw measured by Simoa often preceded these imaging studies.

**Histochemistry: ORF1p immunohistochemistry** was performed as described using anti-ORF1 4H1 (Millipore)(6) diluted 1:3000 on a Leica Bond system. Cases were scored by three experienced gastrointestinal pathologists (MST, VD, OHY). **LINE-1 in situ hybridization** was performed as described using RNAscope catalog 565098 (Advanced Cell Diagnostics) on a Leica Bond system(7). The probe is complementary to the 5' end of L1RP (L1 insertion in X-linked retinitis pigmentosa locus). Cases were scored by three experienced gastrointestinal pathologists (MST, VD, OHY).

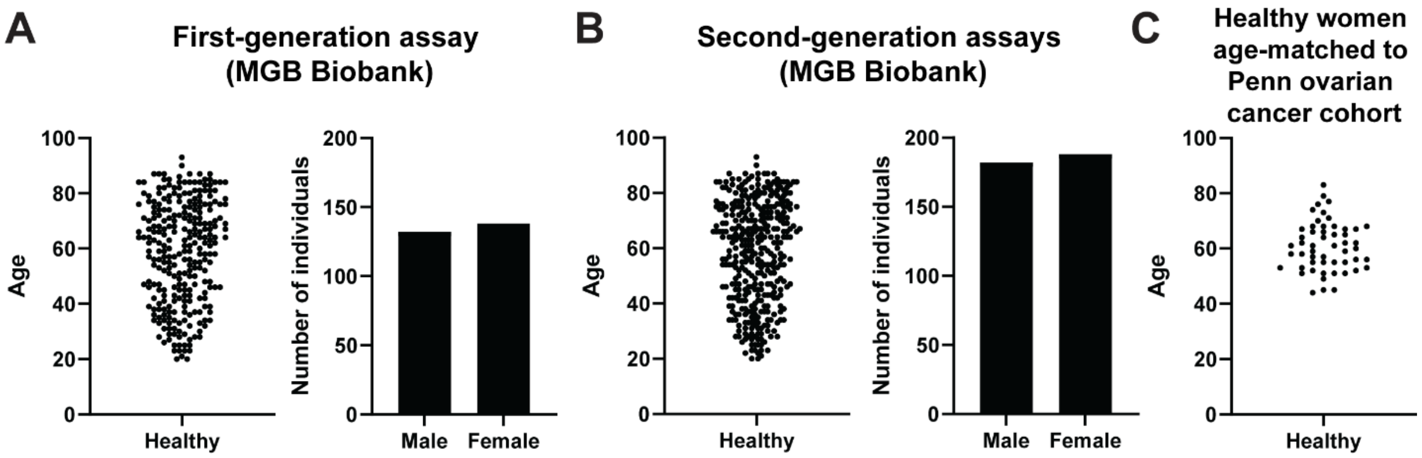


**Supplementary Figure 1.** Age and gender distributions of healthy control patients used in the first- and second-generation ORF1p Simoa assays from the (A-B) Mass General Brigham Biobank in the first-generation (A) and second-generation (B) ORF1p Simoa assays; and (C) Penn Medicine Biobank. Healthy female control patients from the Penn Medicine Biobank were age-matched with the ovarian cancer patients from the Penn Ovarian Cancer Research Center Tumor BioTrust Collection.

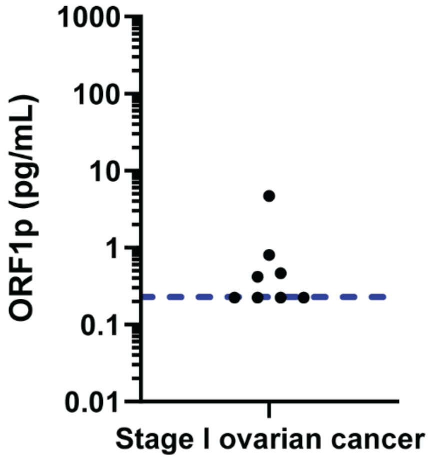


**Supplementary Figure 2.** Circulating ORF1p levels in eight stage I ovarian cancer patients, as measured by the first-generation Simoa assay. Blue dashed line indicates the assay limit of detection, accounting for the four-fold dilution factor.

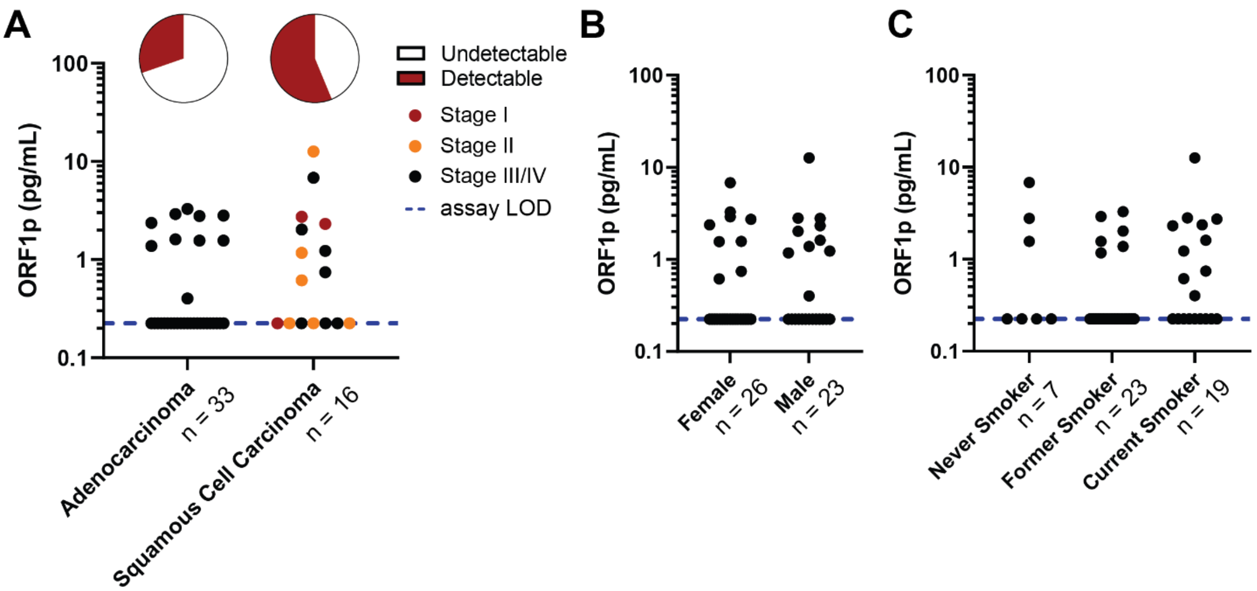


**Supplementary Figure 3.** Circulating ORF1p levels in lung cancer patient cohort, as classified by (A) disease subtype, (B) gender, and (C) smoking status. Blue dashed line indicates the assay limit of detection, accounting for the four-fold dilution factor.

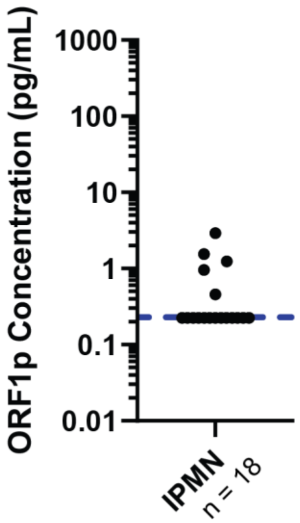


**Supplementary Figure 4.** Circulating ORF1p levels in patients with IPMN, measured using the first-generation Simoa assay. IPMN, intraductal papillary mucinous neoplasm. Blue dashed line denotes the assay limit of detection, accounting for four-fold dilution.

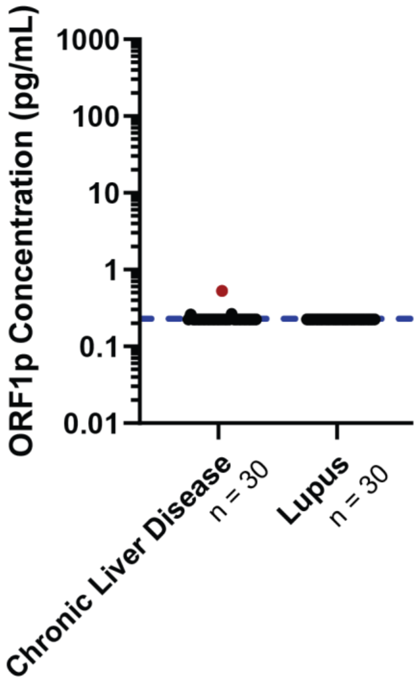


**Supplementary Figure 5.** Circulating ORF1p levels in patients with chronic liver disease or systemic lupus erythematous, measured using the first-generation Simoa assay. The red dot represents a chronic liver disease patient diagnosed with hepatocellular carcinoma after the time of sampling. Blue dashed line denotes the assay limit of detection, accounting for four-fold dilution.

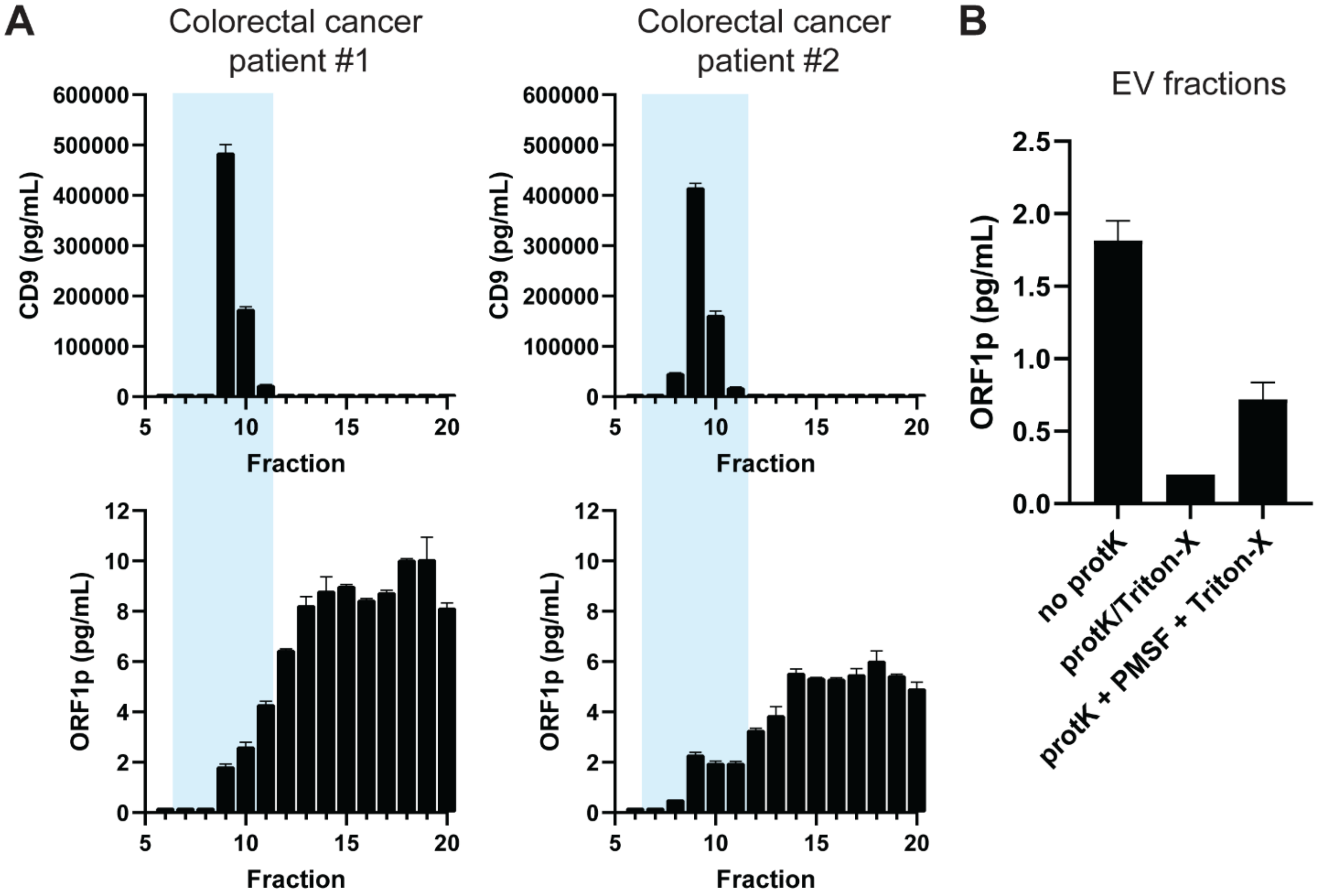


**Supplementary Figure 6.** Analysis of circulating ORF1p levels in size exclusion chromatography (SEC) fractions of colorectal cancer patient plasma. 500 µL plasma was filtered through a 0.45 µm centrifugal filter and fractionated with a Sepharose CL-6B resin packed column (A) CD9 and ORF1p levels in each SEC fraction as measured by Simoa. Blue highlighted boxes denote extracellular vesicle (EV)-containing fractions. The majority of circulating ORF1p is measured in free protein fractions. (B) Determination of ORF1p levels inside EVs via proteinase K (protK) protection assays. EV-containing fractions were pooled and concentrated and treated with proteinase K, followed by the serine protease inhibitor phenylmethylsulfonyl fluoride (PMSF) and 0.5% Triton-X. Controls without proteinase K digestion (no protK) and with simultaneous proteinase K and Triton-X treatment (protK/Triton-X) were performed. Decreased ORF1p concentrations were measured after proteinase K digestion and subsequent EV lysis by Triton-X, further suggesting that only a very small amount of circulating ORF1p is in EVs.

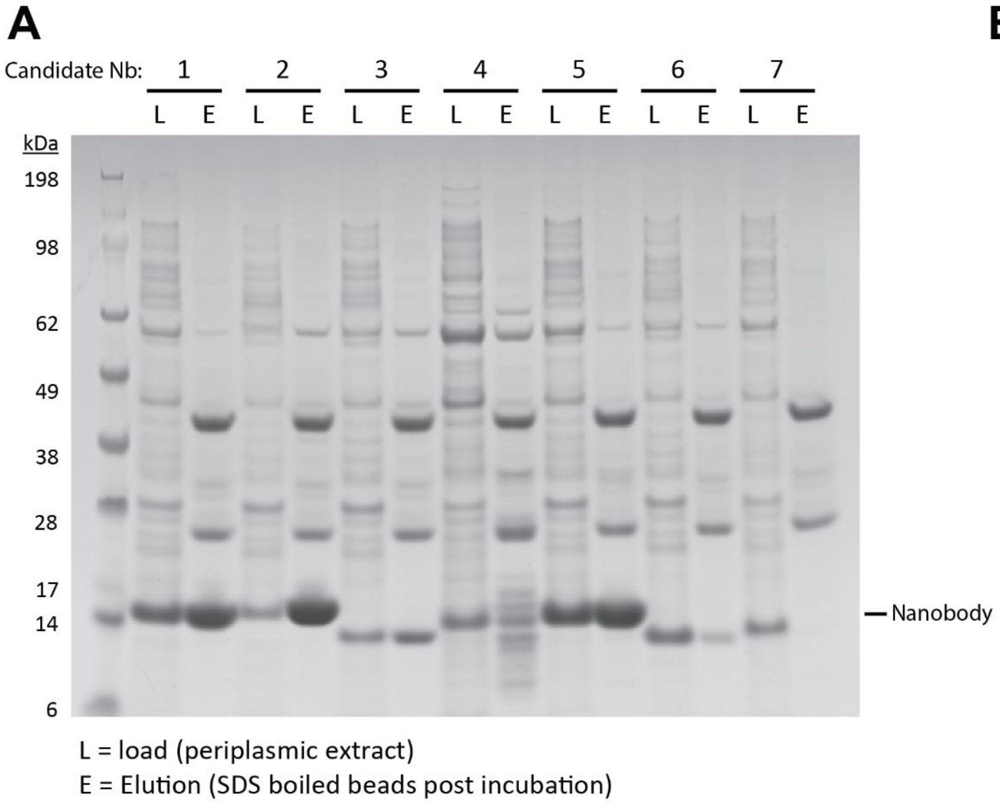


B

| | ELISA EC50 (nM) |
|---|---|
| **Nb ORF1-1** | **19.4** |
| **Nb ORF1-2** | **4.8** |
| Nb ORF1-3 | 3.0 |
| Nb ORF1-4 | 6.0 |
| **Nb ORF1-5** | **1.8** |
| Nb ORF1-6 | 11.4 |
| Nb ORF1-7 | NA |
| **Nb ORF1-8** | **1.4** |
| **Nb ORF1-9** | **1.2** |
| **Nb ORF1-10** | **1.2** |
| **Nb ORF1-11** | **1.5** |
| **Nb ORF1-12** | **1.4** |
| **Nb ORF1-13** | **1.1** |
| Nb ORF1-14 | NA |
| **Nb ORF1-15** | **0.6** |
| **Nb ORF1-16** | **1.0** |
| **Nb ORF1-17** | **1.0** |
| Nb ORF1-18 | 5.6 |
| Nb ORF1-19 | 7.9 |
| **Nb ORF1-20** | **1.8** |
| **Nb ORF1-21** | **0.6** |

**Supplementary Figure 7.** Nanobody generation. **(**A) Representative screening results for nanobodies 1-7. Periplasmic extracts from E. Coli expressing the indicated candidate nanobodies (Nb) were bound to ORF1p-conjugated beads and eluted, then analyzed by SDS-PAGE and stained by Coomassie blue. Expressed and purified nanobodies (~15 kDa) are indicated; larger/darker bands indicate higher levels of expression and/or purification. (B) EC50s from ELISAs performed against recombinant ORF1p. Highlighted nanobodies were selected for follow-up characterization, while remainder were rejected for poor affinity and/or protein expression.

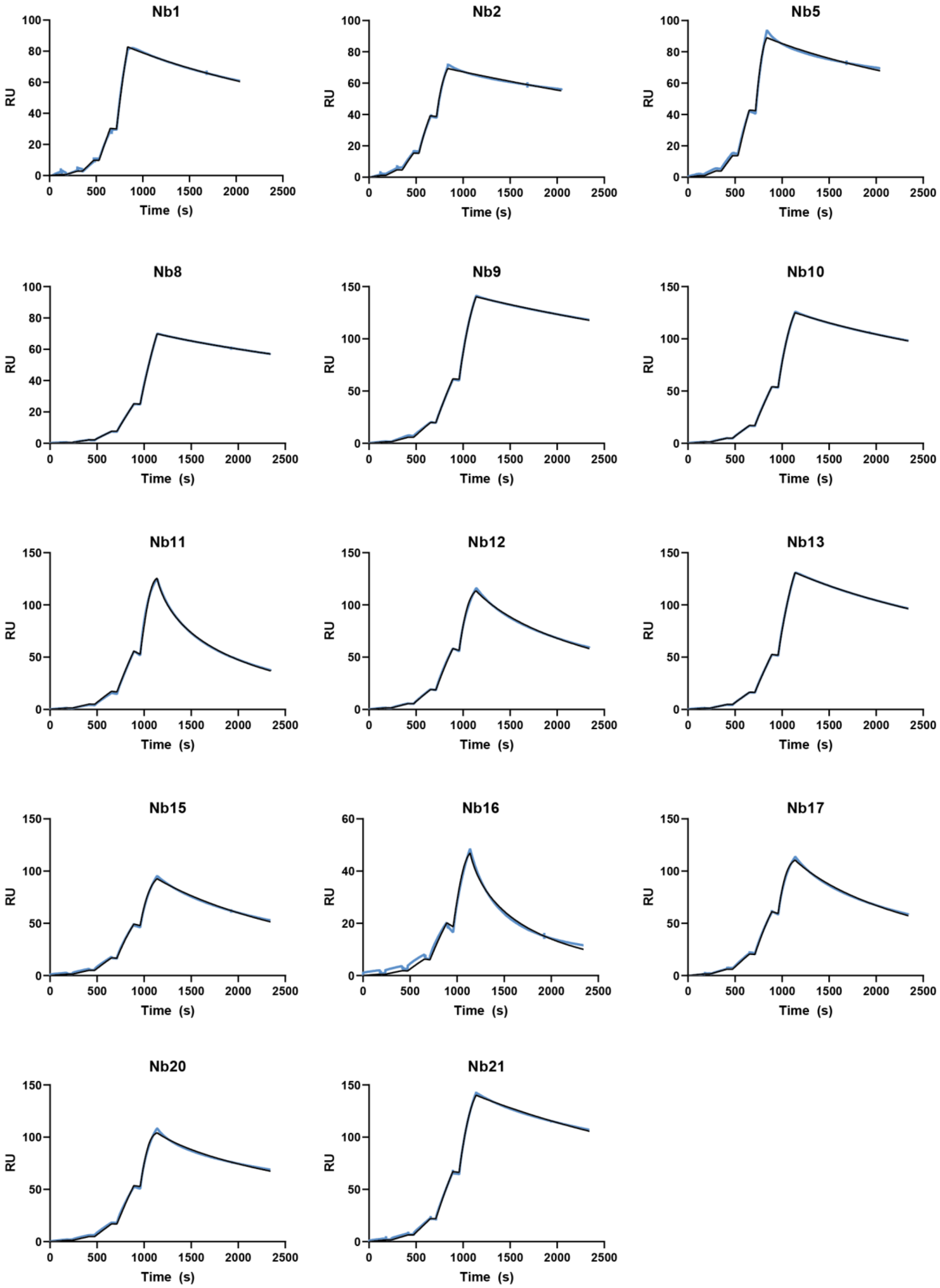


**Supplementary Figure 8.** Representative SPR sensorgrams of ORF1 nanobodies. Single-cycle kinetics performed over immobilized ORF1, with nanobodies injected sequentially at 0.1, 0.3, 1, 3, and 10nM concentrations. Raw data (blue) and a 1:1 binding model fit (black) are plotted.

**A**

**Nanobody Constructs**

| Construct | Description |
|---|---|
| MT991 | Nb 1-1 |
| MT992 | Nb 2-2 |
| MT993 | Nb 5-5 |
| MT994 | Nb 5-1 |
| MT995 | Nb 5-2 |
| MT996 | Nb 5-5-5 |
| MT997 | Nb 5-5 long “LL” |
| MT1032 | Nb 5-5-cys (short) |
| MT1033 | 5-5-cys (long1) |
| MT1034 | 2-2-cys (long2) |
| MT1035 | 5-5-cys(long2) |
| MT1036 | 2-9-cys(long2) |
| MT1037 | 5-9-cys(long2) |
| MT1038 | 9-9-cys(long2) |
| MT1039 | 5-2-9-cys |
| MT1040 | 5-9-2-cys |

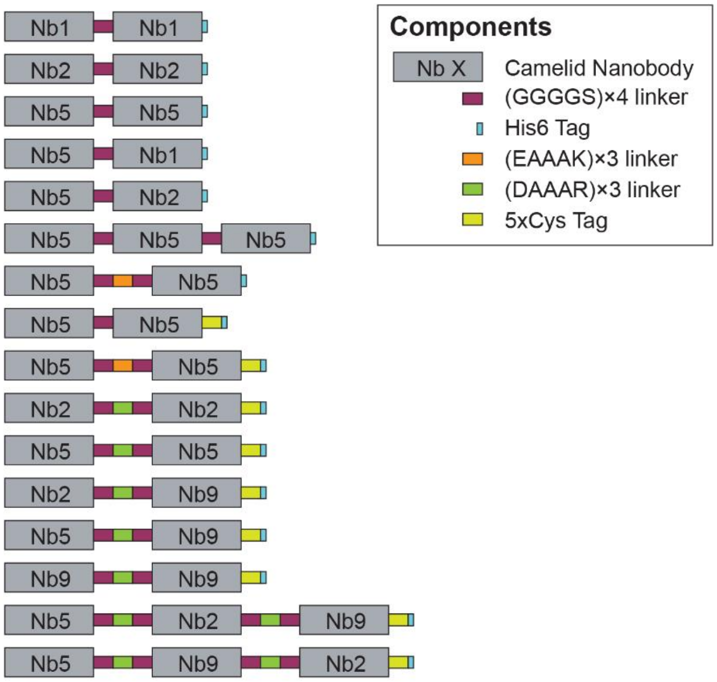


**B**

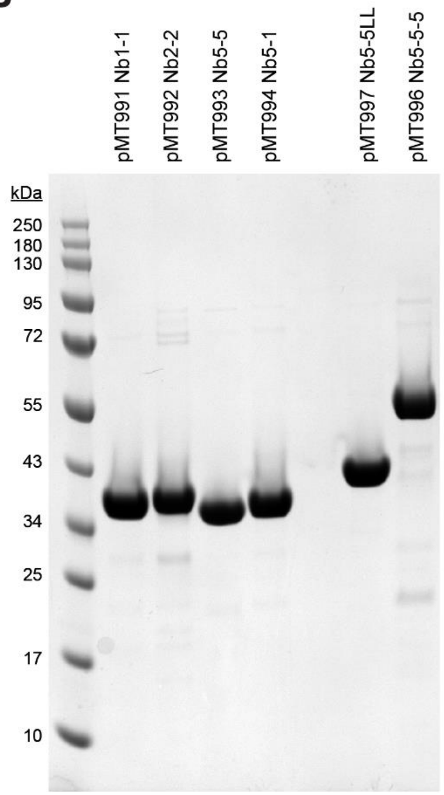


**Supplementary Figure 9. Engineered Nanobody constructs.** (A) Schematic of design of engineered dimeric and trimeric nanobody constructs, with flexible (GGGGS × 4) and rigid helical (EAAAK × 3 or DAAAR × 3) linker. 5xCys tag sequence is CGSGRCGSGRCGSGRCGSGRC. (B), Representative preparation of engineered nanobody constructs, Coomassie stain.

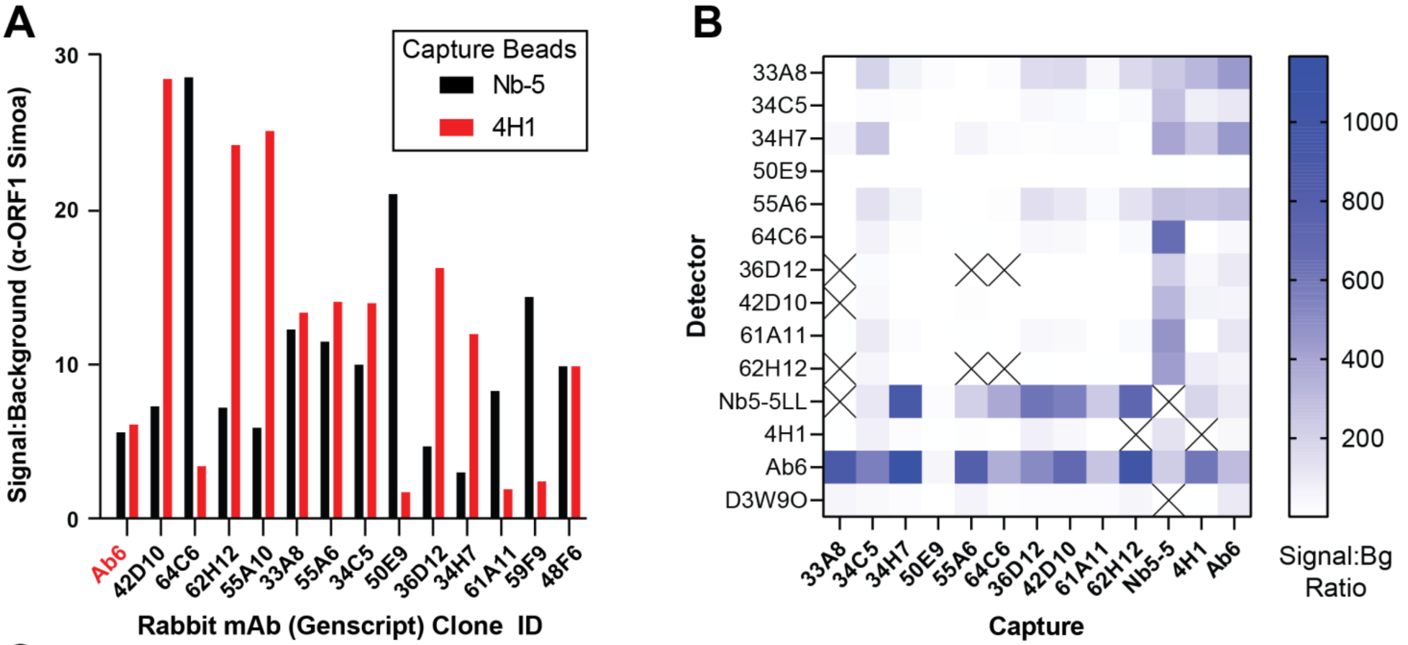


**C**

| | | DETECTOR | | | | | | | | |
|---|---|---|---|---|---|---|---|---|---|---|
| | | **Ab6** | **Nb5-5LL** | **33A8** | **34C5** | **34H7** | **64C6** | **61A11** | **62H12** | **55A6** |
| **CAPTURE** | **33A8** | 0.016 | | | | | | | | |
| | **34C5** | 0.026 | | | | 0.032 | | | | |
| | **34H7** | 0.006 | 0.043 | | | | | | | |
| | **55A6** | 0.026 | 0.055 | | | | | | | |
| | **64C6** | 0.034 | 0.041 | | | | | | | |
| | **36D12** | 0.046 | 0.050 | | | | | | | |
| | **42D10** | 0.021 | 0.020 | | | | | | | |
| | **61A11** | 0.042 | 0.066 | | | | | | | |
| | **62H12** | 0.008 | 0.062 | | | | | | | |
| | **Nb5-5** | 0.050 | | 0.133 | 0.022 | 0.047 | 0.038 | 0.042 | 0.120 | |
| | **4H1** | 0.030 | 0.056 | 0.057 | | 0.044 | | | | 0.036 |
| | **Ab6** | 0.102 | 0.044 | 0.063 | | | | | 0.113 | |

**Limit of Detection (pg/mL)**

**Supplementary Figure 10.** Screening of newly developed monoclonal antibody candidates (GenScript). (A) Signal:background ratios of novel rabbit monoclonal α-ORF1p antibodies (B-cell supernatants) in a modified Simoa employing the candidate mAb plus biotinylated secondary anti-Rabbit pAb as detector; these show up to 5-fold improvement vs. our prior best detection antibody, Ab6. Two different capture beads with distinct epitopes were employed. (B-C) Best performers were then synthesized and purified; mAbs were used in further screening with a dimeric nanobody and commercially available monoclonal antibodies for ORF1p detection with Simoa. (B) Signal-to-background comparisons of affinity reagents as capture/detector pairs on Simoa, using recombinant ORF1p protein. All labeled affinity reagents except Nb5-5LL are monoclonal antibodies; Nb5-5LL denotes a homodimeric form of the nanobody Nb5. (C) Limit of detection values for affinity reagent pairs selected from screening in (B). Anti-ORF1 D3W9O is from Cell Signaling Technology.

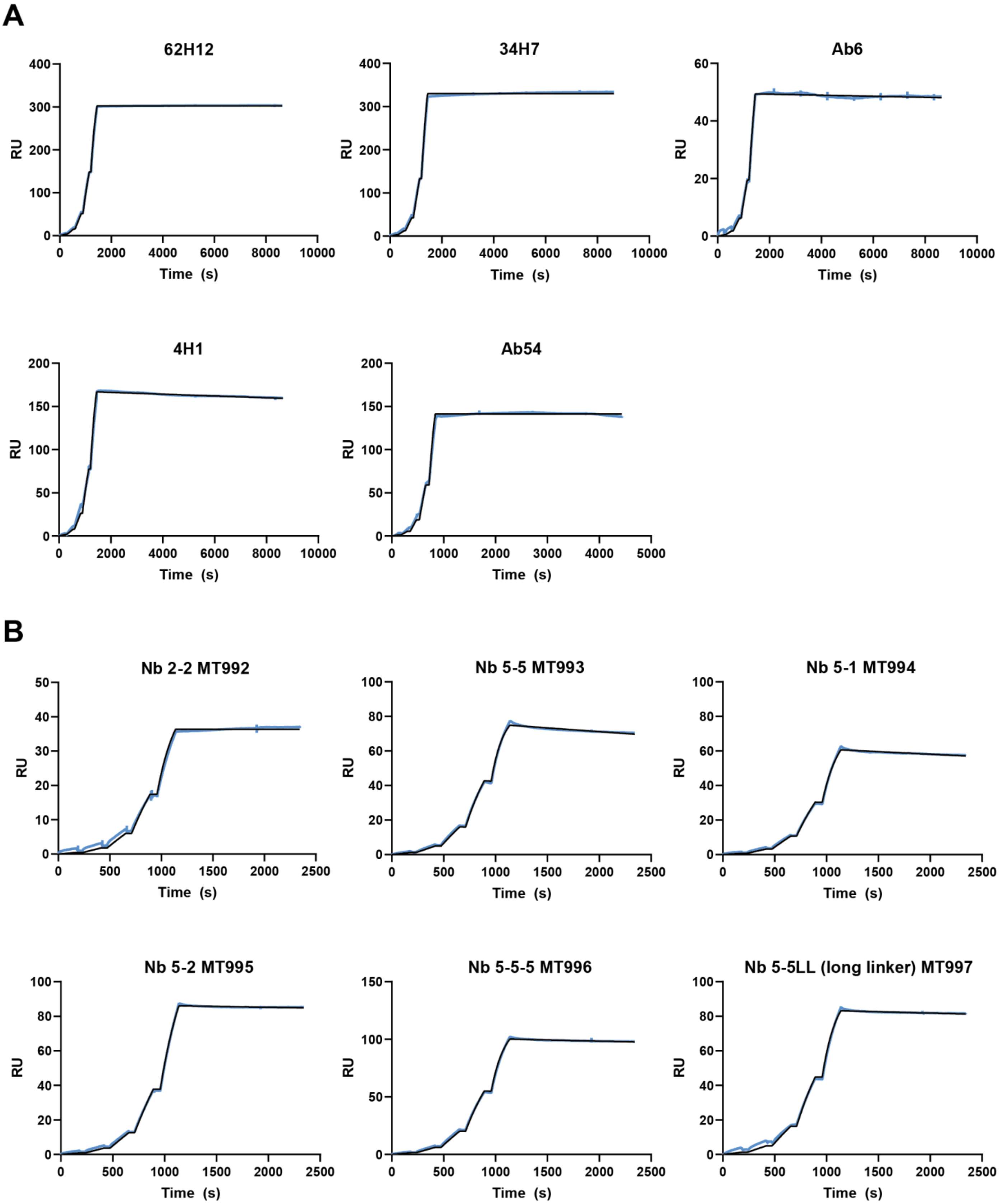


**Supplementary Figure 11.** Representative SPR sensorgrams of ORF1 monoclonal antibodies and nanobody multimers. Single-cycle kinetics performed over immobilized ORF1, with (A) monoclonal antibodies or (B) nanobody multimers injected sequentially at 0.1, 0.3, 1, 3, and 10nM concentrations. Raw data (blue) and a 1:1 binding model fit (black) are plotted.

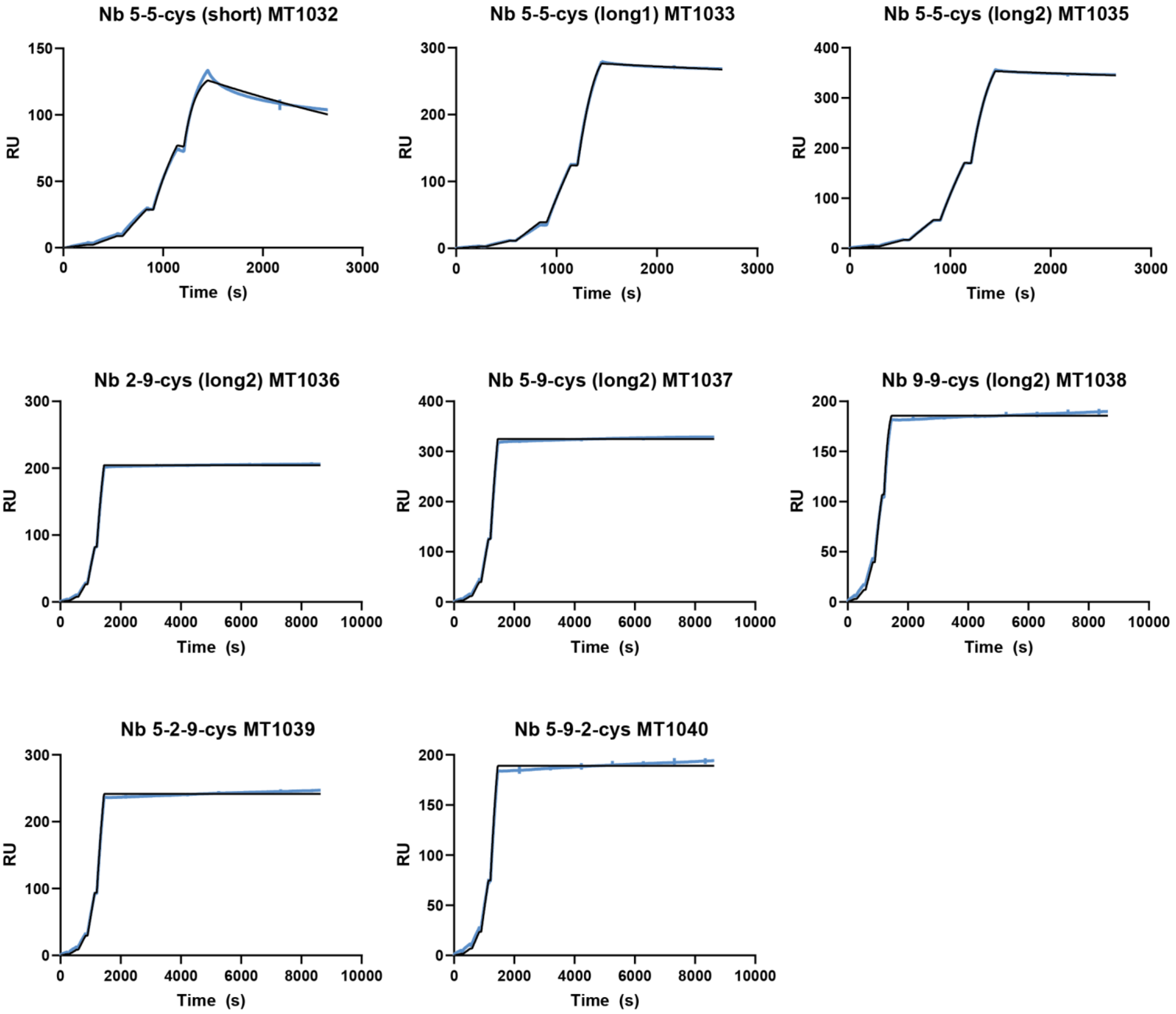


**Supplementary Figure 12.** Representative SPR sensorgrams of ORF1 nanobody multimers with "5xCys Tail". Single-cycle kinetics performed over immobilized ORF1, with nanobodies injected sequentially at 0.1, 0.3, 1, 3, and 10nM concentrations. Raw data (blue) and a 1:1 binding model fit (black) are plotted.

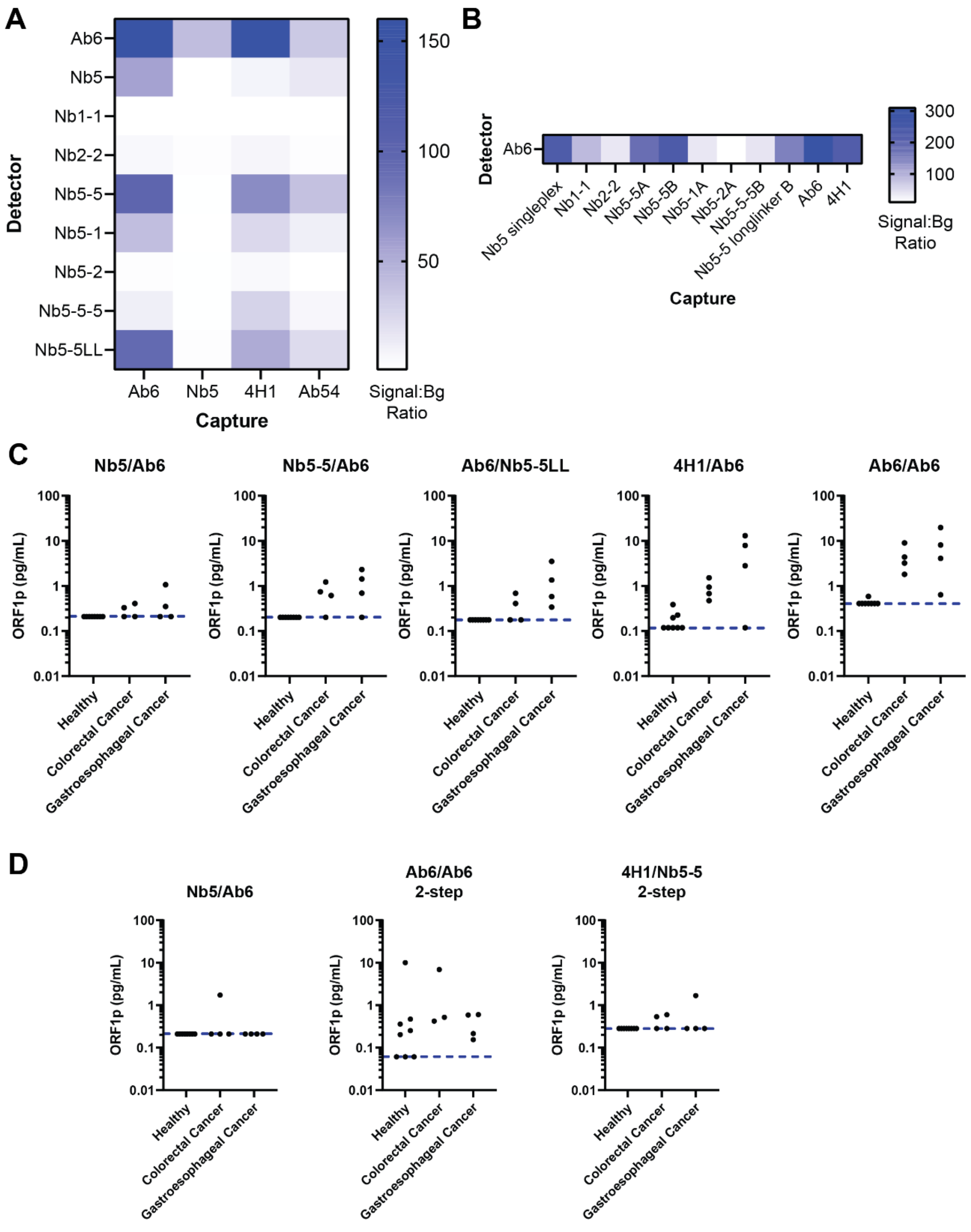


**Supplementary Figure 13.** Screening of multimeric nanobodies for ORF1p detection with Simoa. (A) Signal-to-background (Signal:Bg) comparisons of multimeric nanobodies as detector reagents on the Simoa platform, using recombinant ORF1p protein. A four-plex assay format, with a unique fluorescent dye-encoded bead type for each of the four capture reagents tested, was used for screening. (B) Signal-to-background comparison of multimeric nanobodies as capture reagents on Simoa, using recombinant ORF1p protein, paired with the monoclonal antibody Ab5 as detector reagent. Sub-labels “A” and “B” refer to different nanobody concentrations used during conjugation to beads. (C-D) Screening of select affinity reagent pairs in small sets of plasma samples from healthy and cancer (colorectal and gastroesophageal) patients. Each assay is denoted by the capture/detector reagent pair. The first-generation assay (Nb5/Ab6) measurements are depicted for comparison. All assays were performed as three-step Simoa assays unless otherwise indicated (two-step). Blue dashed lines indicate the assay limits of detection, accounting for four-fold dilution.

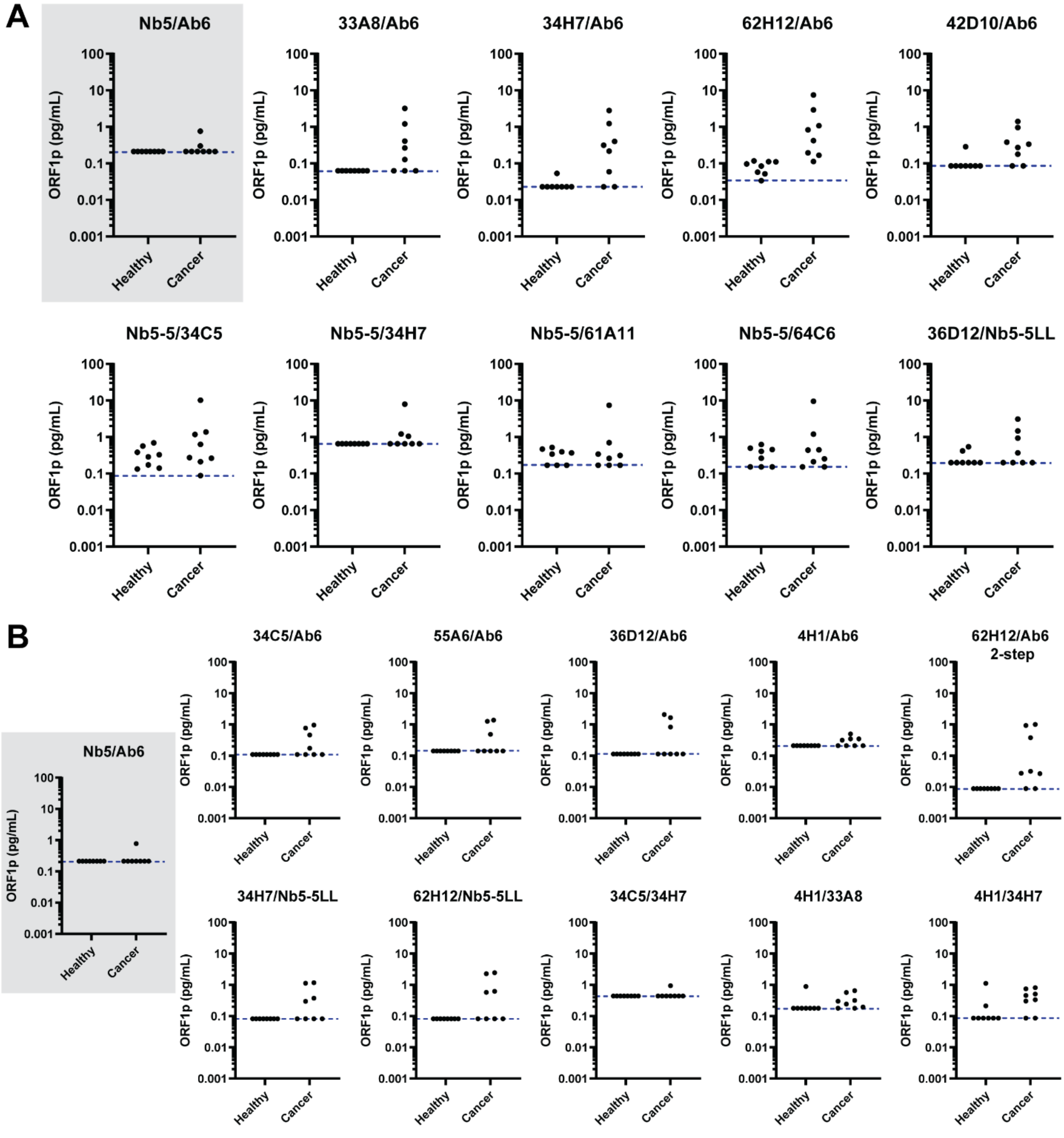


**Supplementary Figure 14.** First round of screening of newly developed monoclonal antibody and dimeric nanobody reagent pairs in patient plasma. Affinity reagent pairs selected from initial screening with recombinant ORF1 protein (Supplementary Figure 8) were screened in small sets of plasma samples from healthy and cancer (colorectal and gastroesophageal) patients. Eight healthy and eight cancer patients were used in each set of assays in (A) and (B). Blue dashed lines indicate assay limits of detection, accounting for four-fold dilution. Each assay is denoted by the capture/detector reagent pair. The first-generation assay (Nb5/Ab6) measurements are depicted for comparison (gray box).

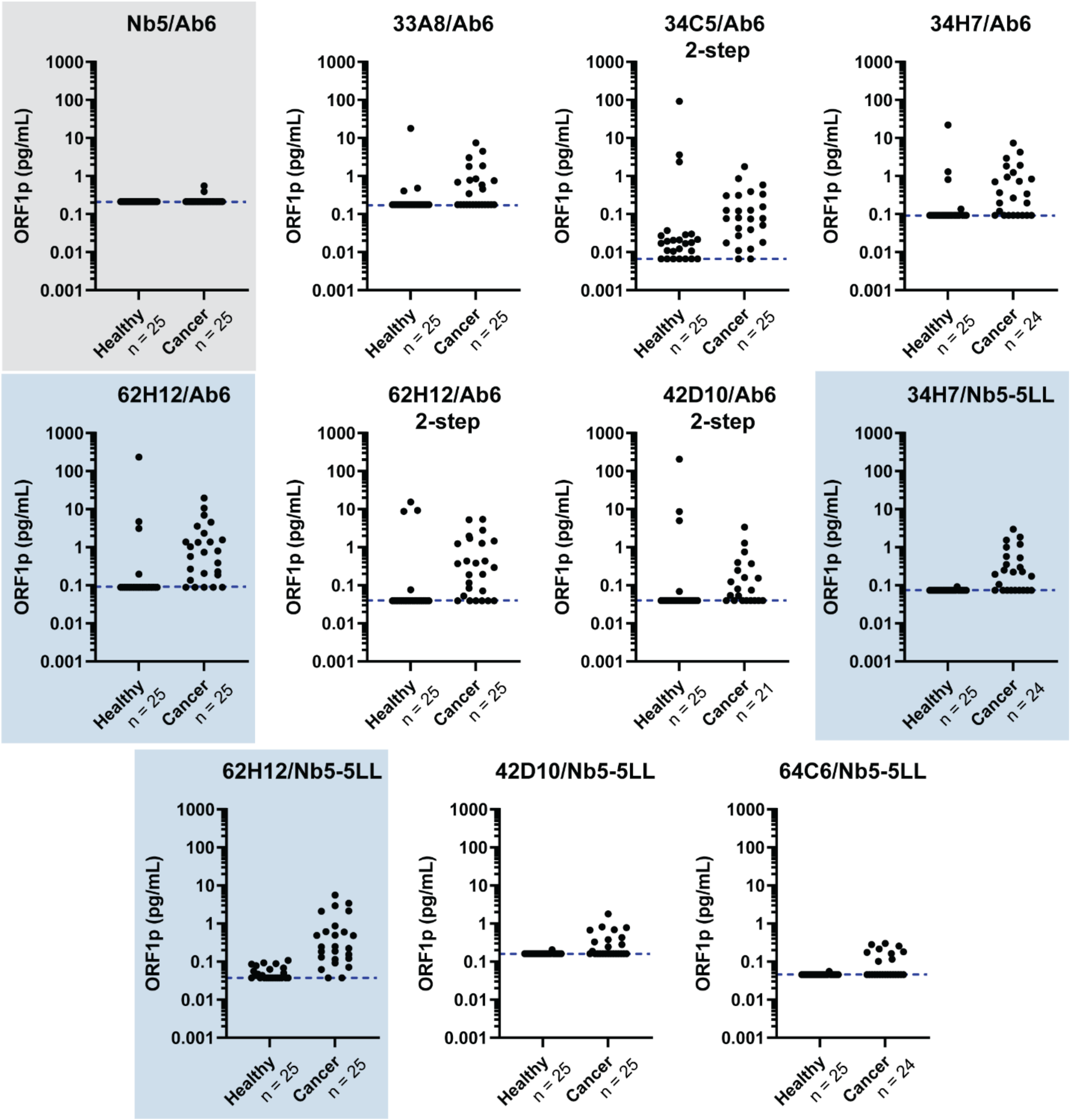


**Supplementary Figure 15.** Second round of screening of newly developed monoclonal antibody and dimeric nanobody reagent pairs in patient plasma. Affinity reagent pairs selected from a first round of screening in plasma samples (Supplementary Figure 9) were screened in 25 healthy and 25 cancer (colorectal, gastroesophageal, and breast) patient plasma samples. Each assay is denoted by the capture/detector reagent pair. The first-generation assay (Nb5/Ab6) measurements are depicted for comparison (gray box). Blue dashed lines indicate assay limits of detection, accounting for four-fold dilution. Blue boxes indicate the final three selected second-generation assays.

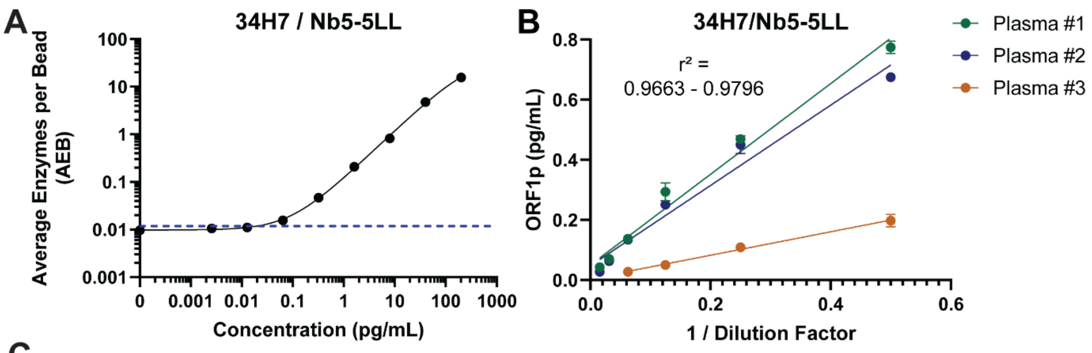


C

| RECOVERY (%) | | | | |
|---|---|---|---|---|
| Spiked Protein (pg/mL) | Plasma #1 | Plasma #2 | Plasma #3 | Plasma #4 |
| 0.2 | 73.8 ± 4.7 | 78.8 ± 9.6 | 67.4 ± 7.9 | 72.1 ± 10.0 |
| 2 | 92.8 ± 0.8 | 82.3 ± 8.2 | 70.7 ± 0.3 | 78.1 ± 5.7 |
| 20 | 83.9 ± 4.3 | 88.4 ± 6.0 | 67.8 ± 6.4 | 77.2 ± 5.8 |

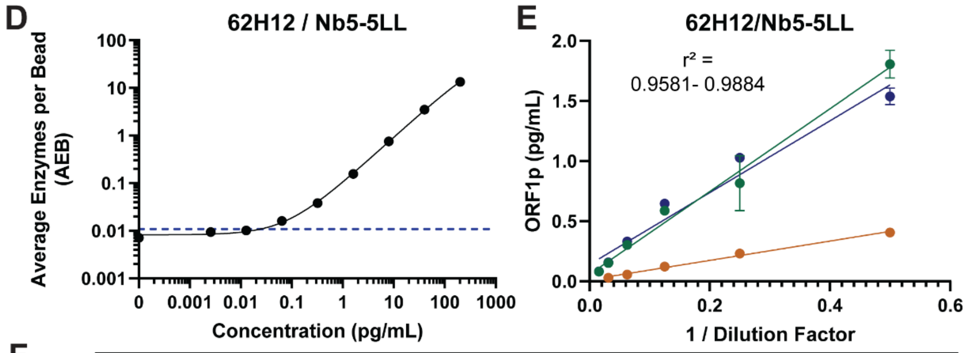


F

| RECOVERY (%) | | | | |
|---|---|---|---|---|
| Spiked Protein (pg/mL) | Plasma #1 | Plasma #2 | Plasma #3 | Plasma #4 |
| 0.2 | 79.1 ± 2.0 | 86.9 ± 9.1 | 64.7 ± 4.9 | 90.5 ± 4.6 |
| 2 | 92.7 ± 2.6 | 88.6 ± 3.2 | 63.9 ± 1.0 | 79.6 ± 1.3 |
| 20 | 103.8 ± 6.4 | 85.4 ± 9.9 | 69.1 ± 5.7 | 84.1 ± 3.2 |

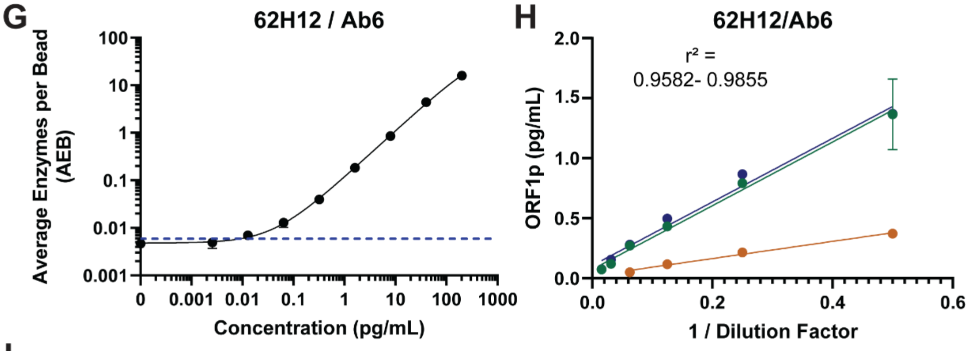


I

| RECOVERY (%) | | | | |
|---|---|---|---|---|
| Spiked Protein (pg/mL) | Plasma #1 | Plasma #2 | Plasma #3 | Plasma #4 |
| 0.2 | 95.8 ± 6.3 | 100.2 ± 0.2 | 84.4 ± 16.7 | 103.4 ± 7.9 |
| 2 | 88.3 ± 6.9 | 97.0 ± 0.9 | 64.7 ± 7.1 | 84.6 ± 0.8 |
| 20 | 85.4 ± 4.2 | 82.2 ± 2.2 | 47.2 | 75.3 ± 1.3 |

**Supplementary Figure 16.** Assay validation for second-generation ORF1p Simoa assays. (A-C) Representative calibration curve (A), dilution linearity (B), and spike and recovery results (C) for the 34H7/Nb5-5LL (capture/detector) assay. (D-F) Representative calibration curve (D), dilution linearity (E), and spike and recovery results (F) for the 62H12/Nb5-5LL (capture/detector) assay. (G-I) Representative calibration curve (G), dilution linearity (H), and spike and recovery results (I) for the 62H12/Ab6 (capture/detector) assay. Error bars represent the standard deviation of triplicate measurements in each calibration curve and duplicate measurements of all plasma samples.

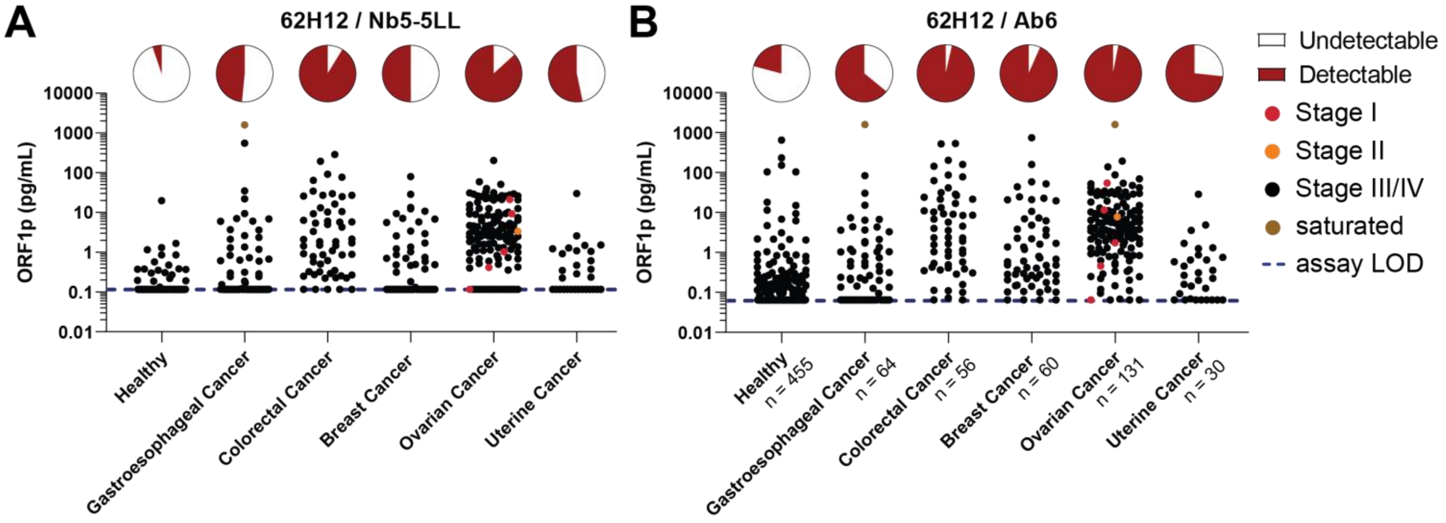


**Supplementary Figure 17.** Additional second-generation ORF1p Simoa assay measurements in healthy and cancer patients. Each assay is denoted by the capture/detector reagent pair. Blue dashed lines indicate assay limits of detection, accounting for four-fold dilution. The proportion of detectable patients within each cancer type is depicted by corresponding pie charts and early-stage patients (stage I/II) are indicated as red and orange dots, respectively.

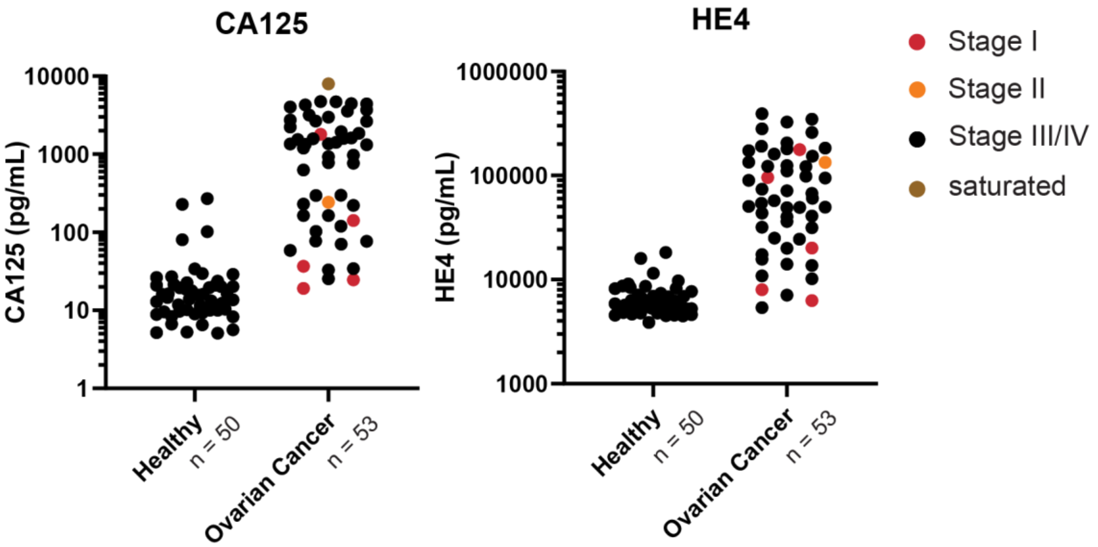


**Supplementary Figure 18.** Circulating CA125 and HE4 levels, as measured by Simoa, in ovarian cancer patient cohort (obtained from Ronny Drapkin at the University of Pennsylvania). Early-stage patients (stage I/II) are indicated as red and orange dots, respectively.

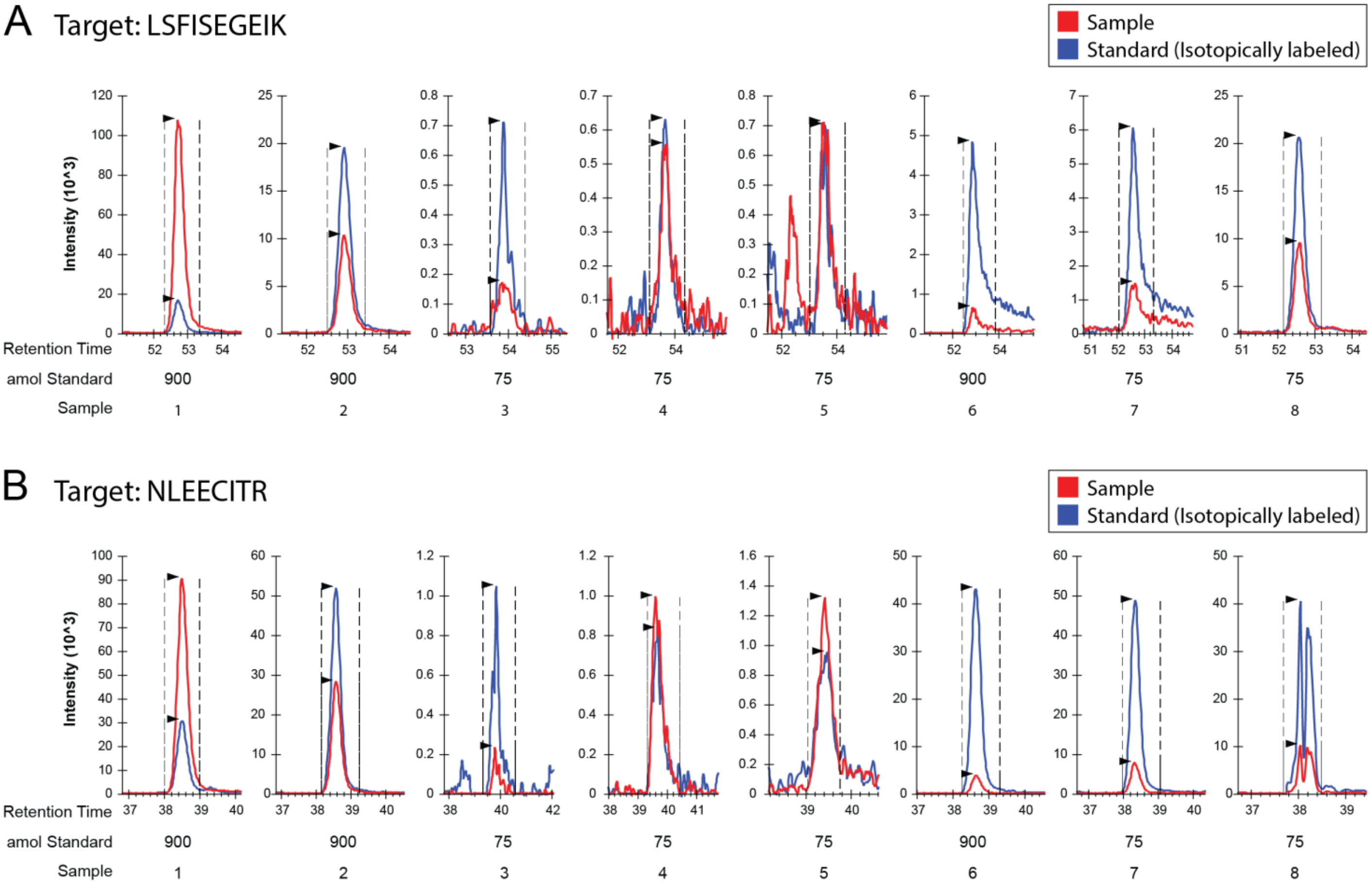


C

| Sample # | Source | Sample Type | ORF1p Simoa (pg/ml) | Plasma Volume (ml) | Total amol Simoa | Total amol LSFISEGEIK | Total amol NLEECIR |
|---|---|---|---|---|---|---|---|
| 1 | MGH | GE Cancer | 1231 | 3.5 | 107444 | 84023 | 36415 |
| 2 | MGH | GE Cancer | 284 | 3 | 21262 | 6345 | 6938 |
| 3 | BioIVT | Healthy | 0.06 | 6 | 9 | 413 | 210 |
| 4 | BioIVT | Pool (2) | 0.24 | 4 | 24 | 1089 | 1210 |
| 5 | BioIVT | Pool (2) | 0.13 | 4 | 13 | 1105 | 1472 |
| 6 | MGH | Healthy | 3.0 | 5 | 374 | 1568 | 1085 |
| 7 | MGH | Healthy | 4.7 | 5 | 586 | 3261 | 2015 |
| 8 | MGH | Healthy | 232 | 5 | 28928 | 5683 | 3370 |

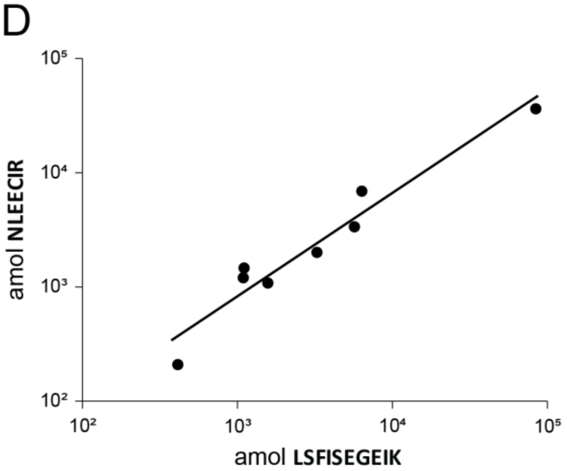


**Supplementary Figure 19.** Targeted proteomics analysis of patient plasma. Samples selected for analysis included two GE cancer patient plasmas with high ORF1p, healthy patients (BioIVT) with very low or undetectable ORF1p, and three healthy patient samples with high ORF1p in the 62H12:Ab6 assay. ORF1p was immunoprecipitated on 62H12-conjugated magnetic beads, digested and prepared for analysis as described above, and assayed for target peptides (A) LSFISEGEIK and (B) NLEECITR, with quantification against spiked-in internal isotopically-labeled standard; results are listed in (C). Traces are from injection of 15% of sample. (C) Assay data: measured and total ORF1p in the samples by Simoa and MS and starting plasma volumes analyzed. Simoa values were from 62H12:Ab6 for samples from MGH (Massachusetts General Hospital) and Nb5:Ab6 for samples from BioIVT; Pool (2): for samples 4 and 5, samples from two healthy patients each were pooled. For healthy samples that were not measurable, the assay LOD was used. (D) Comparison of the two peptides assays. Sample values strongly correlate ($R^2$ = 0.989, linear regression, 99.4% correlation (t test), $p<0.0001$ for both).

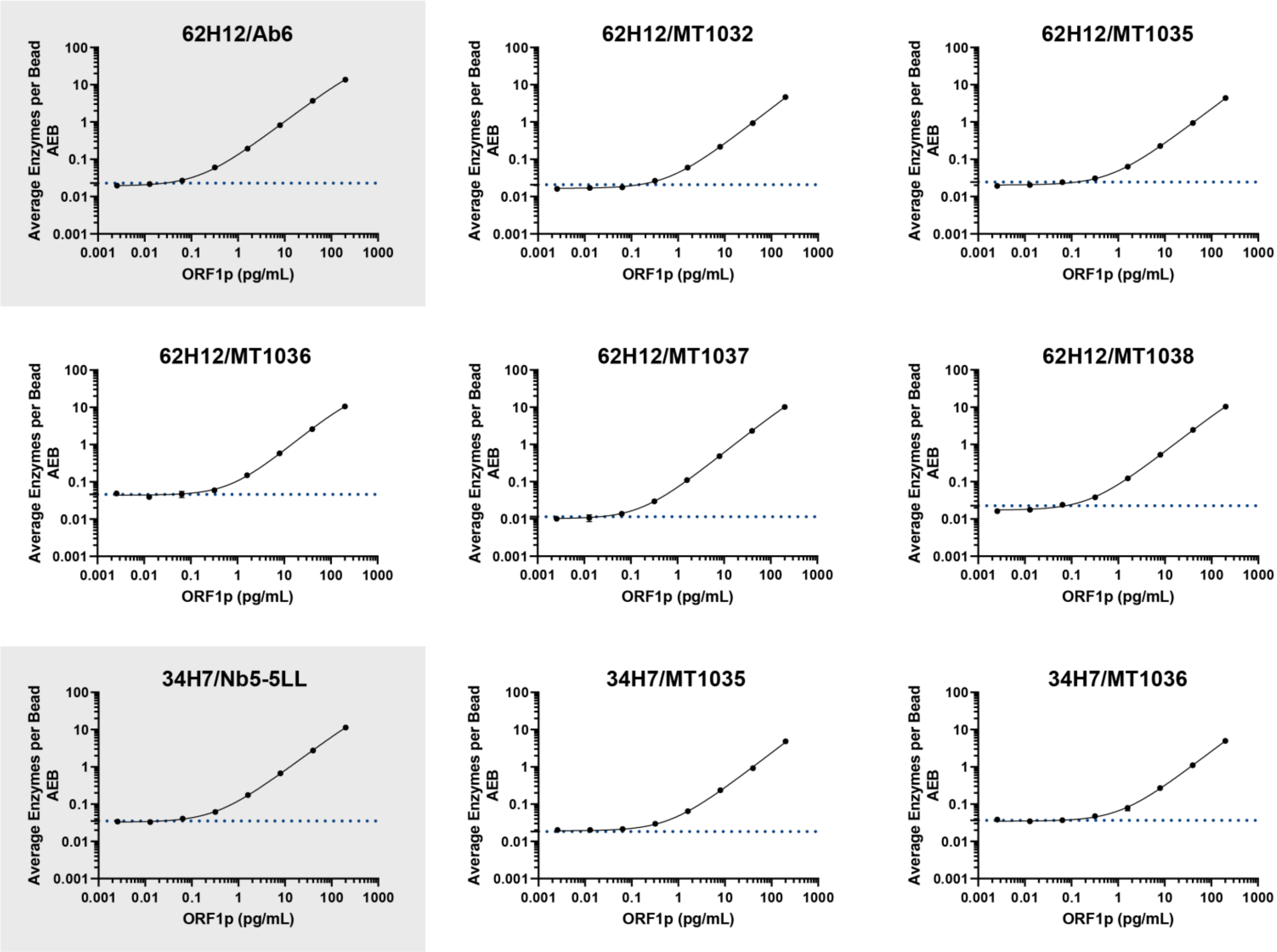


**Supplementary Figure 20.** Calibration curves for “3rd Generation” Simoa assays. Two “2nd Generation” assays are compared, gray boxes. Blue dashed line indicates the assay limit of detection.

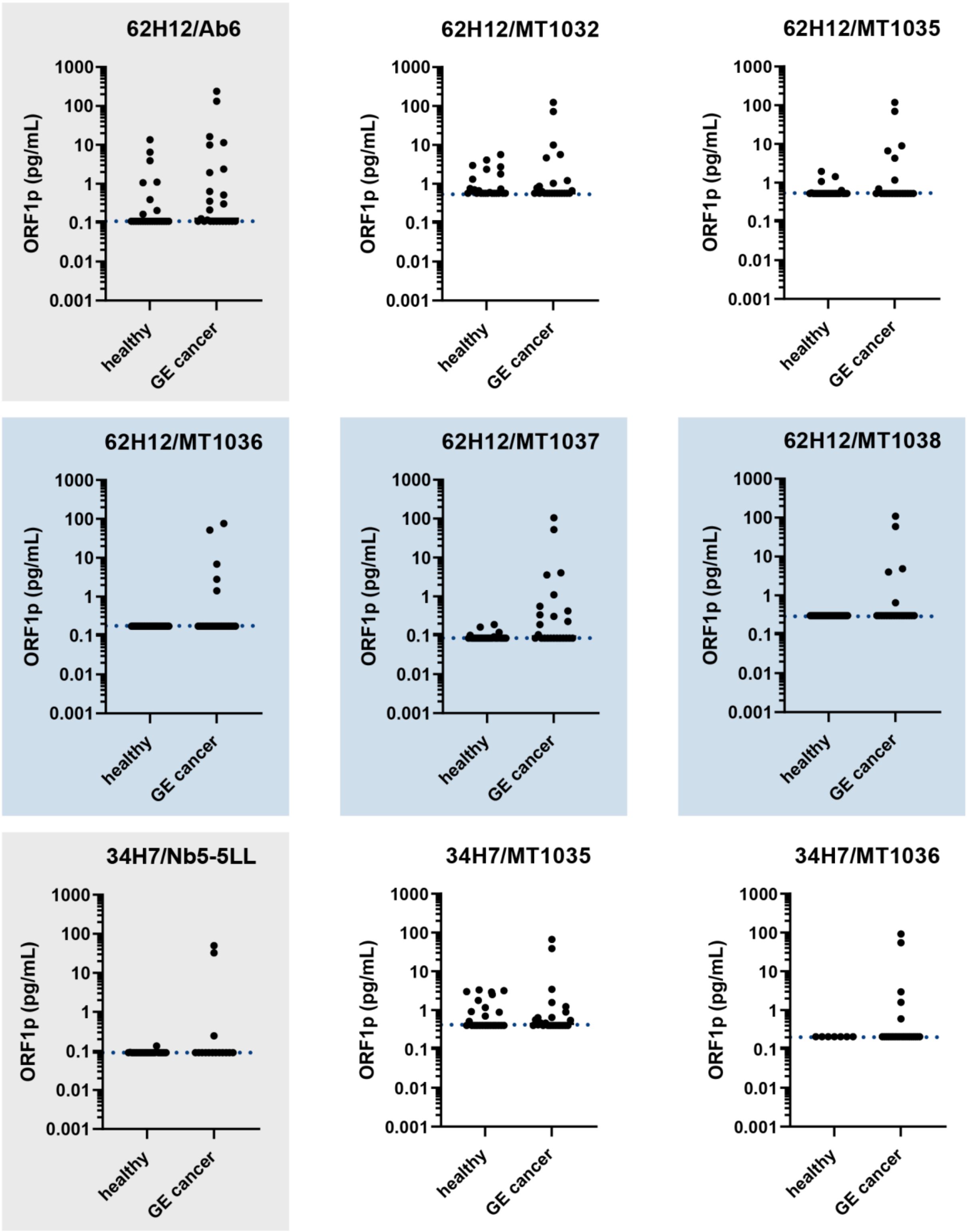


**Supplementary Figure 21.** Second round of screening of newly developed "3rd generation" assays using dimeric nanobody detectors. Affinity reagent pairs selected from a first round of screening in plasma samples (Supplementary Figure 9) were screened in 25 healthy and 25 GE cancer patient plasma samples. Each assay is denoted by the capture/detector reagent pair. Second-generation assays are run for comparison (gray boxes). Blue dashed lines indicate assay limits of detection, accounting for four-fold dilution. Blue boxes indicate the final three selected third-generation assays. MT1032 and MT1035 are Nb5-5 homodimers with different linkers. MT1036 is a Nb2-Nb9 heterodimer, MT1037 a Nb5-Nb9 heterodimer, and MT1038 a Nb9-Nb9 homodimer.

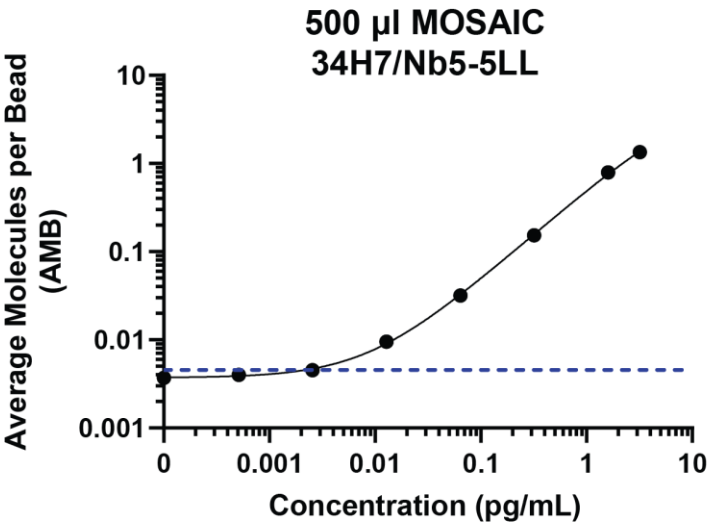


**Supplementary Figure 22.** Calibration curve for the large-volume MOSAIC assay used in Figure 2. Blue dashed line indicates the assay limit of detection.

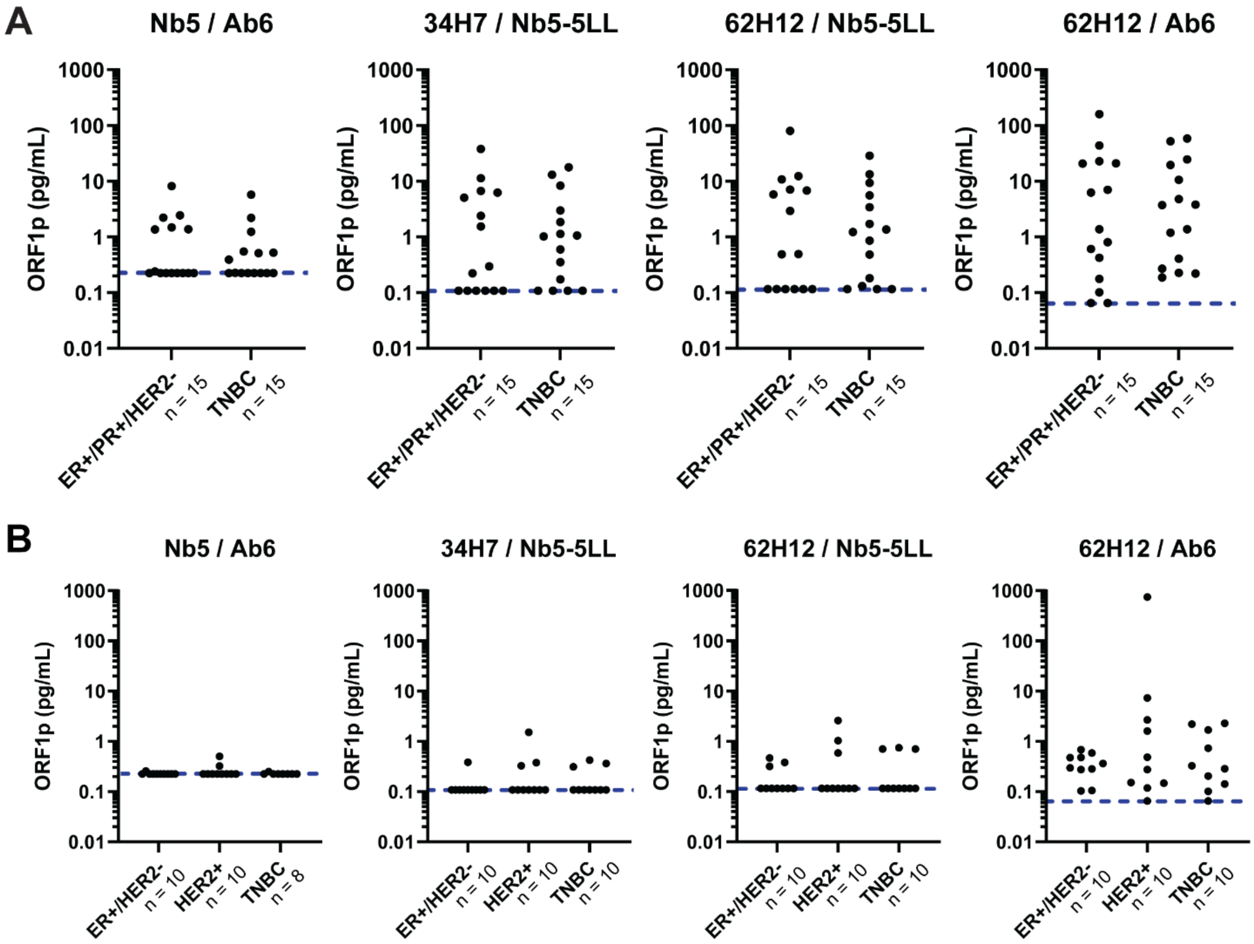


**Supplementary Figure 23.** Circulating ORF1p levels in breast cancer patients, classified by receptor status, with (A) metastatic and (B) localized disease. First- and second-generation Simoa assays were used to measure circulating ORF1p, with each assay labeled as capture/detector reagent. Blue dashed lines indicate assay limits of detection, accounting for four-fold dilution. ER, estrogen receptor; PR, progesterone receptor; HER2, human epidermal growth factor receptor 2; TNBC, triple-negative breast cancer.

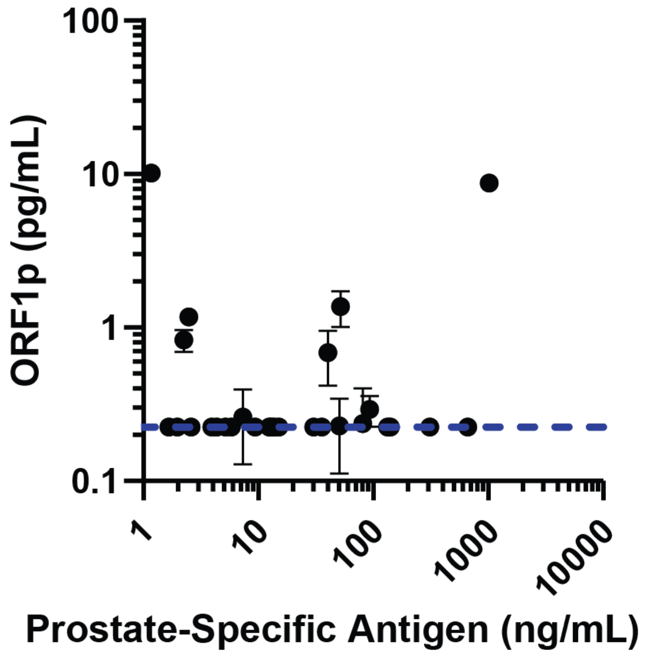


**Supplementary Figure 24.** Circulating ORF1p (first generation assay) versus prostate-specific antigen levels in prostate cancer patient plasma. Blue dashed line indicates the assay limit of detection, accounting for four-fold dilution.

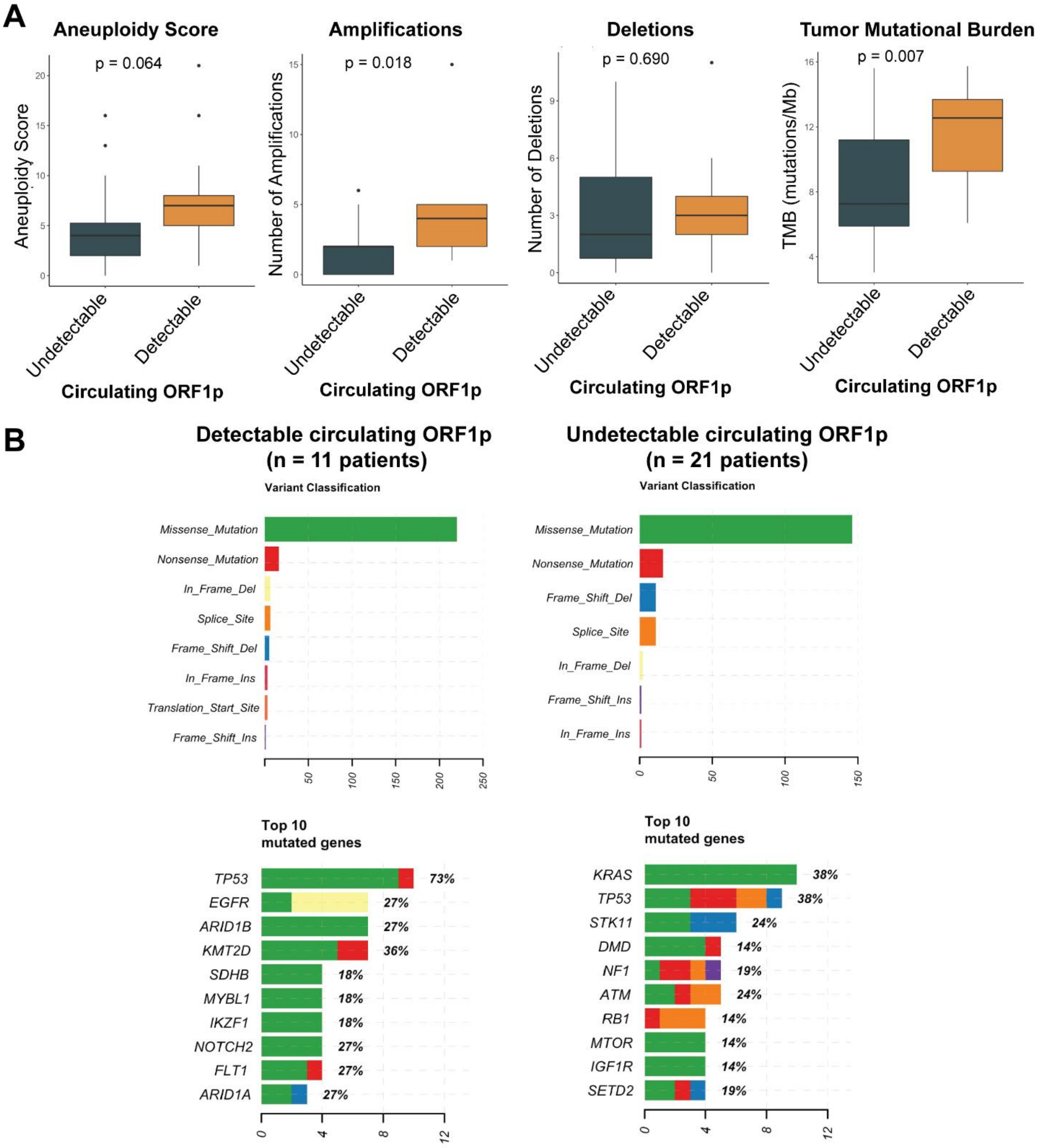


**Supplementary Figure 25.** Genomics analysis of lung cancer patient cohort. (A) Comparisons of aeuploidy score, amplifications, deletions, and tumor mutational burden between lung cancer patients with detectable versus undetectable circulating ORF1p levels, as measured by the first-generation ORF1p Simoa assay. (B) Mutation types and top mutated genes in lung cancer patients with detectable versus undetectable circulating ORF1p.

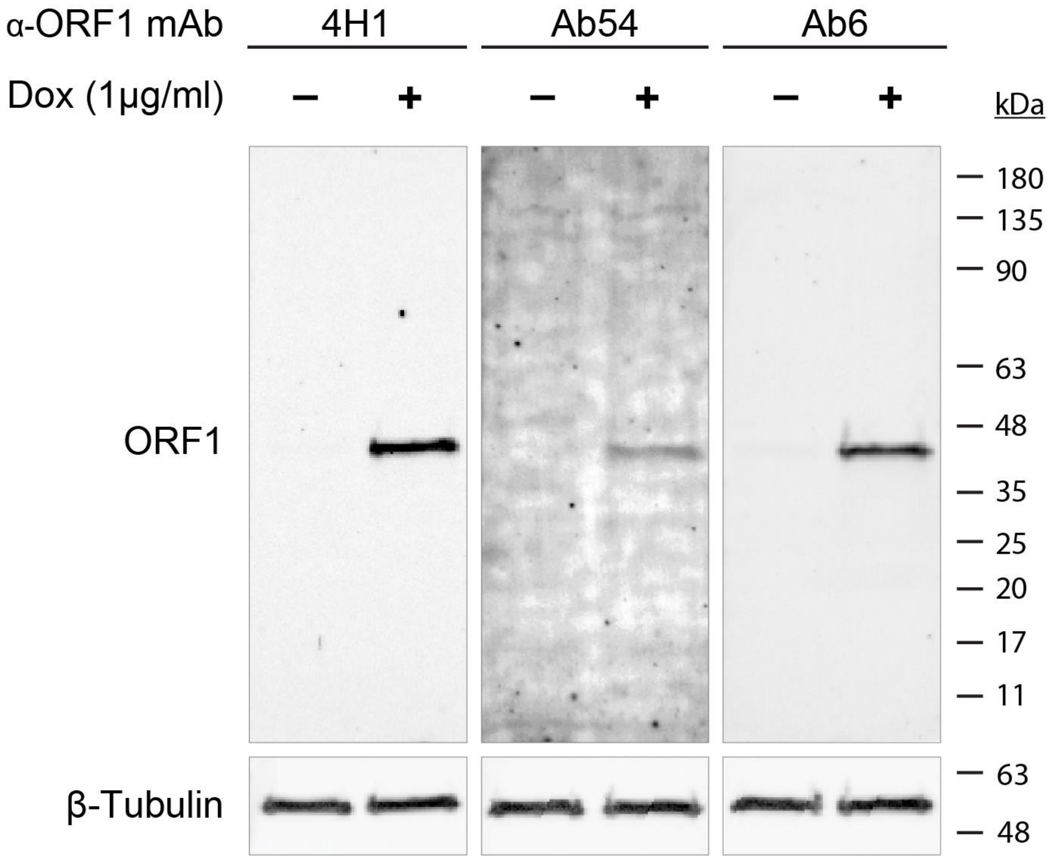


**Supplementary Figure 26.** Validation of monoclonal antibodies by western blotting. HeLa cells with stably integrated tet-On LINE-1 (L1RP)(8) were treated with DMSO as control or with 1 µg/ml doxycycline (Dox) for 4 days to induce L1 expression, lysed with RIPA buffer, cleared, and 20 µg total protein (BCA) was loaded per lane and blotted with the indicated antibodies.

**Table S1.** Assay conditions and limits of detection for first- and second-generation ORF1p assays.

| Platform | Assay (Capture / Detector) | Sample Volume (μL) | Incubation times (capture-detector-streptavidin minutes) | Detector (pg/mL) | Streptavidin β-galactosidase (pM) | Limit of Detection (pg/mL) |
|---|---|---|---|---|---|---|
| Simoa | Nb5 / Ab6 | 100 | 15-5-5 | 0.3 | 150 | 0.056 ± 0.032 |
| Simoa | 34H7 / Nb5-5LL | 100 | 15-5-5 | 0.3 | 300 | 0.027 ± 0.016 |
| Simoa | 62H12 / Nb5-5LL | 100 | 15-5-5 | 0.3 | 300 | 0.029 ± 0.016 |
| Simoa | 62H12 / Ab6 | 100 | 15-5-5 | 0.3 | 300 | 0.016 ± 0.007 |
| MOSAIC | 34H7 / Nb5-5LL | 2000 | 120-10-10 | 0.3 | 100 (streptavidin-DNA) | 0.002 |

**Table S2.** Affinity reagents used across all screening experiments.

| Affinity reagent | Type | Source |
|---|---|---|
| Ab6 | Rabbit monoclonal antibody | Abcam (ab246317) |
| Ab54 | Rabbit monoclonal antibody | Abcam (ab246320) |
| Nb5 | Nanobody | Rockefeller University |
| 4H1 | Mouse monoclonal antibody | MilliporeSigma (MABC1152) |
| JH73 | Rabbit monoclonal antibody | |
| JH74 | Rabbit monoclonal antibody | |
| D3W9O | Rabbit monoclonal antibody | Cell Signaling Technology (88701) |
| Nb5-5 pMT993 | Homodimeric nanobody (short linker) | This study |
| Nb5-5LL pMT997 | Homodimeric nanobody (long linker) | This study |
| Nb1-1 pMT991 | Homodimeric nanobody (short linker) | This study |
| Nb2-2 pMT992 | Homodimeric nanobody (short linker) | This study |
| Nb5-1 pMT994 | Heterodimeric nanobody (short linker) | This study |
| Nb5-2 pMT995 | Heterodimeric nanobody (short linker) | This study |
| 33A8 | Rabbit monoclonal antibody | This study |
| 34C5 | Rabbit monoclonal antibody | This study |
| 34H7 | Rabbit monoclonal antibody | This study |
| 36D12 | Rabbit monoclonal antibody | This study |
| 42D10 | Rabbit monoclonal antibody | This study |
| 50E9 | Rabbit monoclonal antibody | This study |
| 55A6 | Rabbit monoclonal antibody | This study |
| 64C6 | Rabbit monoclonal antibody | This study |
| 61A11 | Rabbit monoclonal antibody | This study |
| 62H12 | Rabbit monoclonal antibody | This study |

**Table S3.** Surface plasmon resonance measurements of binding kinetics of anti-ORF1p nanobodies

| Construct | ka (1/Ms) | kd (1/s) | KD (M) | KD se | n |
|---|---|---|---|---|---|
| NB Orf1-1 | 6.1E+05 | 6.4E-04 | 1.0E-09 | | 1 |
| NB Orf1-2 | 6.4E+05 | 4.2E-05 | 7.1E-11 | 2.0E-11 | 3 |
| NB Orf1-5 | 4.8E+06 | 2.3E-03 | 3.7E-10 | 1.9E-10 | 3 |
| NB Orf1-8 | 9.4E+05 | 2.0E-04 | 2.0E-10 | 4.1E-11 | 3 |
| NB Orf1-9 | 6.1E+05 | 1.3E-04 | 2.2E-10 | 2.0E-11 | 3 |
| NB Orf1-10 | 8.0E+05 | 1.8E-04 | 2.7E-10 | 4.5E-11 | 3 |
| NB Orf1-11 | 4.5E+06 | 3.3E-03 | 6.4E-10 | 1.6E-10 | 3 |
| NB Orf1-12 | 1.2E+06 | 4.2E-04 | 2.9E-10 | 7.6E-11 | 3 |
| NB Orf1-13 | 8.2E+05 | 2.5E-04 | 3.0E-10 | 2.3E-11 | 3 |
| NB Orf1-15 | 1.1E+06 | 5.1E-04 | 4.9E-10 | 3.8E-11 | 3 |
| NB Orf1-16 | 2.6E+06 | 3.2E-03 | 8.4E-10 | 3.4E-10 | 3 |
| NB Orf1-17 | 1.5E+06 | 4.7E-04 | 2.9E-10 | 5.7E-11 | 3 |
| NB Orf1-20 | 1.7E+06 | 3.8E-04 | 2.4E-10 | 4.2E-11 | 3 |
| NB Orf1-21 | 9.6E+05 | 1.8E-04 | 1.7E-10 | 3.3E-11 | 3 |

**Table S4.** Surface plasmon resonance measurements of binding kinetics of affinity reagents used in single molecule assays.

| Construct | # | $k_a$ (1/M·s) | $k_d$ (1/s) | $K_D$ (M) | $K_D$ sem | n |
|---|---|---|---|---|---|---|
| **Dimeric and Trimeric Nanobodies** | | | | | | |
| Nb 1-1 | MT991 | Weak signal | | | | |
| Nb 2-2 | MT992 | 6.4E+05 | <5.0E-06 | <7.9E-12 | 7.4E-13 | 3 |
| Nb 5-5 | MT993 | 9.0E+05 | 3.4E-05 | 3.8E-11 | 1.5E-11 | 3 |
| Nb 5-1 | MT994 | 5.2E+05 | 3.1E-05 | 6.0E-11 | 3.8E-11 | 2 |
| Nb 5-2 | MT995 | 4.0E+05 | 1.3E-05 | 3.4E-11 | 4.7E-12 | 3 |
| Nb 5-5-5 | MT996 | 8.1E+05 | 2.9E-05 | 3.6E-11 | 8.5E-12 | 3 |
| Nb 5-5 long "LL" | MT997 | 7.5E+05 | 2.1E-05 | 2.8E-11 | 1.2E-11 | 3 |
| **"5xCys Tail" Dimeric and Trimeric Nanobodies** | | | | | | |
| Nb 5-5-cys (short) | MT1032 | 1.2E+06 | 1.7E-04 | 1.5E-10 | 4.7E-11 | 3 |
| 5-5-cys (long1) | MT1033 | 1.1E+06 | 6.0E-05 | 5.2E-11 | 2.5E-11 | 3 |
| 2-2-cys (long2) | MT1034 | Weak signal | | | | |
| 5-5-cys(long2) | MT1035 | 8.5E+05 | 3.9E-05 | 4.5E-11 | 1.7E-11 | 3 |
| 2-9-cys(long2) | MT1036 | 9.0E+05 | <5.0E-06 | <5.6E-12 | 3.6E-12 | 3 |
| 5-9-cys(long2) | MT1037 | 5.9E+05 | <5.0E-06 | <8.4E-12 | 3.6E-12 | 3 |
| 9-9-cys(long2) | MT1038 | 1.1E+06 | <5.0E-06 | <4.6E-12 | 1.8E-12 | 4 |
| 5-2-9-cys | MT1039 | 7.5E+05 | <5.0E-06 | <6.7E-12 | 4.5E-12 | 3 |
| 5-9-2-cys | MT1040 | 3.9E+05 | <5.0E-06 | <1.4E-11 | 2.7E-12 | 4 |
| **Monoclonal Antibodies** | | | | | | |
| mAb | 62H12 | 4.8E+05 | <5.0E-06 | <1.0E-11 | 4.4E-13 | 3 |
| mAb | 34H7 | 5.2E+05 | <5.0E-06 | <9.6E-12 | 4.1E-12 | 3 |
| mAb | Ab6 | 2.2E+05 | <5.0E-06 | <2.2E-11 | 2.6E-12 | 3 |
| mAb | 4H1 | 4.5E+05 | <5.0E-06 | <1.1E-11 | 2.5E-12 | 3 |
| mAb | Ab54 | 4.2E+05 | <5.0E-06 | <1.2E-11 | 5.5E-14 | 2 |

**Table S5.** Epitope binning of ORF1p nanobodies. Fraction of 2nd antibody able to bind after saturation with 1st antibody is indicated, normalized to maximum signal bound. Overlapping epitopes are defined by <0.3 remaining binding activity (red shading). Epitope groups are labeled I-III.

| | | **2nd Ab** | | | | | |
|---|---|---|---|---|---|---|---|
| | **Group:** | **I** | **II** | **III** | | | |
| | | **Nb-2** | **Nb-5** | **Nb-8** | **Nb-9** | **Nb-10** | **Nb-21** |
| **1st Ab** | **Nb-2** | 0.16 | 0.66 | 0.94 | 1.00 | 1.00 | 1.00 |
| | **Nb-5** | 0.82 | -0.06 | 1.00 | 0.99 | 0.93 | 0.90 |
| | **Nb-8** | 1.00 | 0.53 | 0.07 | 0.13 | 0.26 | 0.24 |
| | **Nb-9** | 0.91 | 0.66 | 0.03 | 0.08 | 0.19 | 0.17 |
| | **Nb-10** | 0.70 | 1.00 | -0.08 | -0.07 | 0.08 | 0.05 |
| | **Nb-21** | 0.83 | 0.47 | -0.06 | -0.05 | 0.08 | 0.03 |

**Table S6**. Epitope binning of ORF1p MAbs and representative nanobodies. Fraction of 2nd antibody able to bind after saturation with 1st antibody is indicated, normalized to maximum signal bound. Overlapping epitopes are defined by <0.3 remaining binding activity (red shading). Epitope groups are labeled I-V.

| | | **2nd Ab** | | | | | |
|---|---|---|---|---|---|---|---|
| | **Group:** | **I** | **II** | **III** | **IV** | **V** | |
| | | **Nb-2** | **Nb-5** | **Nb-9** | **Ab6** | **34H7** | **62H12** |
| **1st Ab** | **Nb-2** | 0.18 | 0.91 | 0.82 | 0.97 | 1.00 | 0.98 |
| | **Nb-5** | 0.67 | 0.02 | 0.83 | 0.95 | 0.88 | 0.99 |
| | **Nb-9** | 0.64 | 0.90 | 0.04 | 0.78 | 0.42 | 0.86 |
| | **Ab6** | 1.00 | 0.94 | 0.87 | 0.07 | 0.71 | 1.00 |
| | **34H7** | 0.87 | 0.92 | 0.85 | 0.81 | 0.10 | 0.22 |
| | **62H12** | 0.89 | 1.00 | 1.00 | 1.00 | 0.59 | 0.12 |

**Table S7.** Performance metrics of classification models built from all healthy and all ovarian cancer patients measured by the second-generation Simoa assays. Classification models were built using five-fold cross validation, with univariate logistic regression for each individual assay and k-nearest neighbors algorithm for the multivariate model combining two assays.

| Assay | Accuracy | Sensitivity | Specificity | F1 Score |
|---|---|---|---|---|
| 34H7 / Nb5-5LL | 95.0% | 85.7% | 97.5% | 0.876 |
| 62H12 / Nb5-5LL | 94.6% | 86.5% | 96.9% | 0.870 |
| 62H12 / Ab6 | 92.0% | 87.3% | 93.3% | 0.827 |
| 34H7/Nb5-5LL and 62H12/Ab6 | 94.4% | 84.1% | 97.3% | 0.865 |

**Table S8.** Performance metrics of classification models built from ovarian cancer and age-matched healthy female patients (Ronny Drapkin, University of Pennsylvania). Classification models were built using k-nearest neighbors (KNN) and light gradient-boosting machine (LightGBM) algorithms, with five-fold cross validation.

| Biomarkers | Accuracy | Sensitivity | Specificity | F1 Score |
|---|---|---|---|---|
| CA125 and HE4 | 89.0% | 90.0% | 88.0% | 0.891 |
| CA125, HE4, and ORF1p (34H7/Nb5-5LL) | 95.0% | 94.0% | 96.0% | 0.949 |
| CA125, HE4, and ORF1p (62H12/Nb5-5LL) | 93.0% | 88.2% | 98.0% | 0.926 |
| CA125, HE4, and ORF1p (62H12/Ab6) | 91.1% | 86.2% | 96.0% | 0.906 |

**Table S9.** Clinicopathological characteristics for gastroesophageal cancer cohort

| Characteristic | | N=19 |
|---|---|---|
| Median age (range)-yr | | 76 (37-81%) |
| Male sex-no. (%) | | 19 (100%) |
| Histology-no. (%) | Adenocarcinoma | 19 (100%) |
| Primary tumor location no. (%) | Esophagus | 7 (37%) |
| | Gastroesophageal Junction | 7 (37%) |
| | Stomach | 5 (26%) |
| Disease stage at initial diagnosis-no. (%) | Locally advanced (II-III) | 11 (58%) |
| | Advanced (IV) | 8 (42%) |
| Treatment paradigm during pre- and post-assay collection* | Neoadjuvant | 9 (47%) |
| | Adjuvant | 2 (11%) |
| | Palliative | 8 (42%) |
| | Surgery | 3 (16%) |
| Treatment response | Responder | 13 (68%) |
| | Non-responder | 6 (32%) |

*Patients who underwent surgery also received systemic therapy thus total exceeds N=19

**Table S10.** Demographic and clinicopathological variables of head and neck squamous cell carcinoma patients grouped by detectability of circulating ORF1p.

| | | Line1 Orf1 Plasma Status | | | |
|---|---|---|---|---|---|
| | | Total (n=50) | Negative (n=38) | Positive (n=12) | p* |
| Age, median (range) | | 64.8 (20.8-87.9) | 65.6 (20.8-87.9) | 62.3 (52.2-83.7) | 0.718 |
| Sex | | | | | |
| | Male | 36 | 26 | 10 | 0.468 |
| | Female | 14 | 12 | 2 | |
| Cancer Site | | | | | |
| | Oral Cavity | 37 | 28 | 9 | 1.000 |
| | Oropharynx | 4 | 3 | 1 | |
| | Larynx/Hypopharynx | 9 | 7 | 2 | |
| Race | | | | | |
| | White | 44 | 33 | 11 | 0.388 |
| | Asian | 2 | 2 | 0 | |
| | Black | 1 | 0 | 1 | |
| | Other/Declined | 3 | 3 | 0 | |
| pT Stage | | | | | |
| | pT2 | 7 | 5 | 2 | 0.623 |
| | pT3 | 13 | 11 | 2 | |
| | pT4a | 21 | 15 | 6 | |
| | Unknown | 9 | 7 | 2 | |
| pN Stage | | | | | |
| | pN0 | 20 | 16 | 4 | 0.111 |
| | pN1 | 8 | 6 | 2 | |
| | pN2 | 6 | 6 | 0 | |
| | pN3 | 7 | 3 | 4 | |
| | Unknown | 9 | 7 | 2 | |
| PNI | | | | | |
| | Positive | 19 | 15 | 4 | 0.731 |
| | Negative | 25 | 18 | 7 | |
| | Unknown | 6 | 5 | 1 | |
| ENE | | | | | |
| | Positive | 9 | 6 | 3 | 1.000 |
| | Negative | 14 | 10 | 4 | |
| | NA (pN0) | 19 | 15 | 4 | |
| | Unknown | 8 | 7 | 1 | |

*Rank sum test p value provided for continuous variable; Fisher's exact test p values provided for categorical data; unknown and NA categories excluded for statistical analyses. PNI, perineural invasion. ENE, extranodal extension.
